# A structural competition involving WDR5 times circadian oscillations

**DOI:** 10.1101/2023.06.05.543739

**Authors:** Arne Börgel, Audrey Rawleigh, Marcel Conrady, Florian Hof, Kyra Ricci, Steven Brown, Eva Wolf

## Abstract

In the mammalian circadian clock, the transcription factor BMAL1/CLOCK cycles between an active state recruiting co-activators like MLL1, and repressed states associated with the clock proteins Period1/2 (PER1/2) and Cryptochrome1/2 (CRY1/2). The MLL1 complex component WDR5 was also found in a repressive PER complex, but the roles of WDR5-PER interactions are unknown. Here we show that WDR5 directly binds to the C-terminal CRY binding domain (CBD) regions of PER1 and PER2. PER2 binds WDR5 via a WDR5 binding (WBM) motif within the PER2/CRY interface, imposing a molecular choice between PER2/WDR5- and PER2/CRY complexes. PER1 predominantly binds WDR5 via a WDR5 interacting (WIN) motif outside the CBD and exhibits a 15-fold higher WDR5 affinity than PER2. Thereby PER1 can form trimeric PER1_WIN_/WDR5/RbBP5_WBM_ - and PER1/WDR5/CRY complexes as potential transition states between activating MLL1_WIN_/WDR5/RbBP5_WBM_ - and repressive PER/CRY complexes. Overexpressing WDR5 in mammalian cells increases the circadian amplitude, whereas a compound targeting the WIN motif binding site of WDR5 weakens PER1-WDR5 interactions and shortens the circadian oscillation period by 2 to 3 hours. Together, our studies uncover WDR5 as direct PER interaction partner at the interface between active- and repressed states of BMAL1/CLOCK and suggest a functional role of WDR5 and its WIN site interactions in the mammalian clock by both enhancing circadian oscillations and creating a temporal delay.

## Main

In mammals, circadian clocks regulate many cellular and physiological processes like cell division, metabolism, sleep-wake cycles and the immune response ^1^. Disruption of circadian rhythms enhances the risk and severity of many diseases including sleep and mood disorders, metabolic syndrome, obesity, cardiovascular diseases, inflammation and also cancer ^2–4^. Circadian clocks are operated by cell-autonomous transcriptional and translational feedback loops. Two transcription factors, *circadian locomotor output cycles kaput* (CLOCK) and *brain and muscle ARNT-like 1* (BMAL1) heterodimerize to activate genes of the clock proteins Period1/2 (PER1/2), Cryptochrome1/2 (CRY1/2), REV-ERBα/β and RORα/γ as well as many clock-controlled genes. During the circadian cycle, BMAL1/CLOCK transitions between transcriptionally active-, early repressed- and late repressed states. In the active state, BMAL1/CLOCK recruits co-activators such as the histone acetyltransferase p300/CBP and the histone methyltransferase *mixed-lineage leukaemia 1* (MLL1). The early repressed state, where CRY and PER repress BMAL1/CLOCK within a large multi-subunit complex, is followed by a late repressed state, where only CRY1 remains bound to BMAL1/CLOCK ^5^. The molecular sequence of events leading from circadian gene activation to repression remains largely unknown but is thought to involve a transition from active (open) to repressed (closed) chromatin ^6^. Timed to arrive at circadian promotors at the interface between activation and repression ^7,8^, PER proteins likely play a key role in the transition process.

Several distinct PER protein complexes have been identified, including large nuclear repressive complexes and two smaller cytosolic complexes ^7,9–13^. Apart from PER, CRY and casein kinase 1δ/ε (CK1δ/ε) these complexes contain RNA binding- and chromatin modifying proteins, including histone deacetylases, repressive methyltransferases, the Drosophila-Behavior-Human-Splicing (DBHS) family members NONO and SFPQ/PSF, and the *WD repeat-containing protein 5* (WDR5) ^5,7,9,10,13^. Within these complexes, PER1 and PER2 are thought to have partially redundant roles but also have distinct impacts on the circadian clock. While disruption of PER1 leads to shorter circadian periods in mice ^14,15^ as well as a lower circadian amplitude and shortened period in cultured cells ^16^, PER2 inactivation results in a loss of circadian rhythm in both systems ^16,17^. Furthermore, PER1 is present on circadian promotors several hours before PER2^7–9^, and PER1 and PER2 are associated with different chromatin modifying complexes ^7^. Both, PER1 and PER2 can dimerize or interact with other proteins via their PAS (PER-ARNT-SIM) domains ^18,19^. Additionally, CK1δ/ε binds to the central PER region ^20^, while CRY1/2 bind to the C-terminal region of PER1 and PER2 (Fig. 1a) ^21–23^.

**Figure 1:**
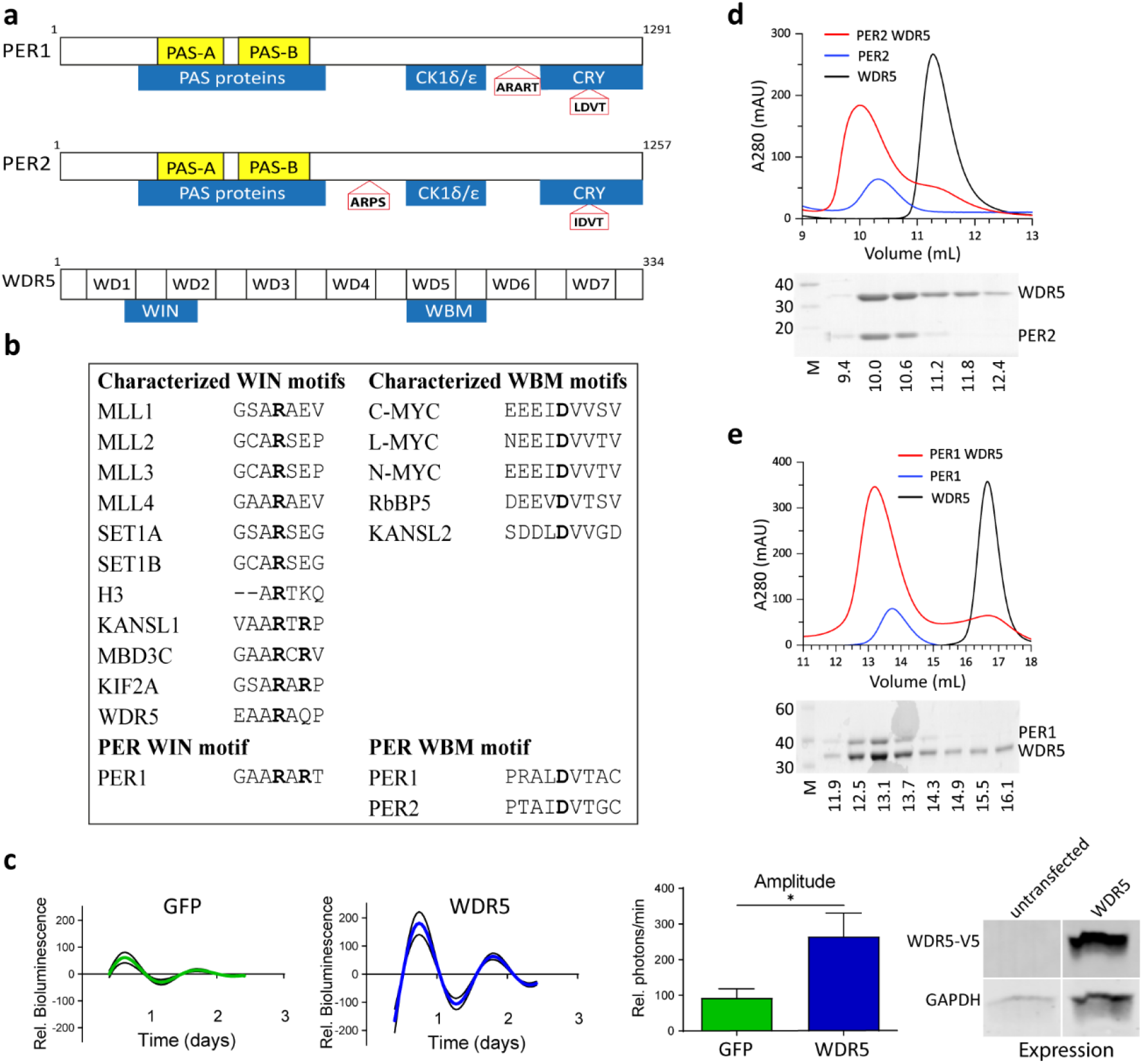
WDR5 directly binds to the CRY-binding regions of PER1 and PER2 and increases the circadian oscillation amplitude. **a** Domains (yellow) and interaction partners (blue) of PER1, PER2 and WDR5. Red boxes indicate potential WDR5 binding WIN (ARART, ARPS) - and WBM (L/IDVT) motifs in PER1/2. PER1/2 have two PAS (PER-ARNT-SIM) domains (PAS-A, PAS-B) to dimerize and interact with other proteins. The middle and C-terminal regions of PER1/2 interact with CK1δ/ε and CRY1/2, respectively. WDR5 consists of 7 WD repeats (WD1-7) and has two binding sites to interact with proteins containing a WIN- or WBM motif. **b** Alignment of WDR5 binding motifs. WIN motifs have a conserved central arginine (bold), WBM motifs consist of a conserved aspartate (bold) flanked by aliphatic amino acids. **c** WDR5 overexpression increases the amplitude of circadian Bmal1-luc oscillations in human U2OS cells. *Left:* Bioluminescence of U2OS cells transiently transfected with the Bmal1-luc circadian reporter construct and either GFP (control) or WDR5 (n = 6 replicates from 3 independent experiments). Data are plotted as an average (colored lines) with the SEM (black lines). *Middle:* relative amplitude (arbitrary units), * P = 0. 035 (two-tailed unpaired t test) shows statistically significant amplitude difference between GFP and WDR5. *Right:* western blot of WDR5-V5 expression and GAPDH loading control in transiently transfected U2OS cells. **d** Analytical SEC (S75 10/300) of WDR5 23-334 and PER2 1132-1252. *Top:* chromatograms of WDR5 23-334 (black), PER2 1132-1252 (blue) and PER2/WDR5 complex (red). *Bottom*: SDS-PAGE of PER2/WDR5 complex SEC. M: molecular weight marker. The PER2/WDR5 complex elutes earlier (10 mL) than free WDR5 (> 11 mL). **e** Analytical SEC (S200 10/300) of WDR5 23-334 and PER1 1013-1291. *Top:* chromatograms of WDR5 23-334 (black), PER1 1013-1291 (blue) and PER1/WDR5 complex (red). *Bottom:* SDS-PAGE of Per1/WDR5 complex SEC. The PER1/WDR5 complex elutes earlier (∼13 mL) than free WDR5 (> 16 mL).

For its part, WDR5 is involved in several cellular processes by interacting with transcription factors (e.g. Myc) and by acting as a scaffold protein in various nuclear multi-protein complexes, such as the SET1/MLL histone H3K4 methyltransferase complexes, the non-specific lethal (NLS) histone acetyltransferase complex or the nucleosome remodeling and deacetylase (NuRD) complex. It possesses two binding pockets that interact with *WDR5 interacting* motifs (= WIN motifs, A/S/C-A-**R-** A/S/C/T) and with *WDR5 binding motifs* (= WBM motifs, -L/I/V-**D**-V-V/T) provided by its varying interaction partners (Fig. 1a,b) ^24^. Within the circadian clock, WDR5 plays an important role as part of the MLL1 complex, which is recruited to BMAL1/CLOCK in a circadian manner to co-activate BMAL1/CLOCK dependent transcription by di- and tri-methylation of histone H3 lysine K4 (H3K4) ^25,26^. Depletion of WDR5 abolishes circadian methylation rhythms of both H3K4 and H3K9, suggesting a role of WDR5 in transcriptional activation (H3K4 methylation) and repression (H3K9 methylation) ^9^. The active MLL1 complex consists of five proteins, namely the *Histone-lysine N-methyltransferase 2A* (KMT2A, MLL1), WDR5, *Retino-blastoma binding protein 5* (RbBP5), A*bsent small homeotic-2-like protein* (Ash2l) and DPY30, and its presence functionally enhances histone methylation activity by several hundredfold ^27^. Within this complex WDR5 binds directly to the WIN motif of MLL1 (3758 GSARAEV 3764) and to the WBM motif of RbBP5 (373 EEVDVT 378) (Fig. 1b) ^28–32^.

While WDR5 is a well characterized MLL1 complex component, its role in repressive PER containing complexes of the mammalian circadian clock is unknown. Moreover, we are far from understanding the protein interaction networks that lead from transcriptional activation to repression of circadian genes at a structural and mechanistic level. To address these open questions and to shed light into the suggested dual role of WDR5 in circadian transcriptional activation and repression, we have analyzed the molecular interactions of PER1 and PER2 with WDR5, and their relationship to repressive CRY- and activating MLL1 complexes. We show that WDR5 directly binds to the C-terminal CRY-binding region of either PER1 or PER2. PER2 binds to the WBM motif binding pocket (= WBM site) of WDR5, resulting in binding competition between CRY and WDR5 on PER2. In contrast, PER1 predominantly binds to the WIN motif binding pocket (= WIN site) of WDR5 and exhibits a 15-fold higher affinity for WDR5 than PER2. Thereby, PER1 can form trimeric PER1/WDR5/RbBP5- and PER1/WDR5/CRY complexes as potential transition states between the active MLL1/WDR5/RbBP5 complex and repressive PER/CRY containing complexes. Functionally, we demonstrate, that this molecular dance both enhances circadian oscillations and creates a temporal delay: whereas overexpression of WDR5 results in an enhanced circadian amplitude, the C6 compound targeting the WDR5 WIN site ^33^ weakens the PER1-WDR5 interaction and significantly shortens the period of circadian oscillations by 2 to 3 h, documenting a functional role of WDR5 and its WIN site interactions in the mammalian circadian clock. Together, our studies uncovered a new role of WDR5 as a direct PER interaction partner in the mammalian circadian clock and revealed critical differences between PER1- and PER2 interactions with WDR5. By uncovering an unexpected strong effect of the C6 WIN site inhibitor on the circadian clock, our results are of high relevance for the ongoing development of WDR5 WIN site inhibitors in the context of MLL1- and MYC-related cancers ^24,34^ and suggest future applications of such compounds as clock modulators and potentially in the treatment of clock related diseases ^2–4^.

## Results

### WDR5 directly interacts with the CRY-binding regions of PER1 and PER2 and affects circadian oscillations

WDR5 was reported to interact with PER1 and PER2 in cell-based Co-IP studies ^9^. Furthermore, using a BMAL1-Luc reporter bioluminescence assay ^35^, we showed that transient overexpression of WDR5 significantly increases the amplitude of endogenous circadian oscillations in human U2OS cells as compared to GFP transfected control cells (Fig. 1c). This indicates that WDR5 is involved in the mammalian circadian clock, potentially enhancing its robustness by increasing the circadian oscillation amplitude. Moreover, the reported Co-IP studies suggest that interactions between PER and WDR5 are important for the clock-related functions of WDR5.

To find out if WDR5 directly interacts with PER and to determine the involved interaction sites, we set out to characterize the WDR5-PER interactions using purified proteins. Interestingly, we found a potential WBM motif within the C-terminal CRY-binding domain (CBD) ^21^ of PER1 (^1196^L**D**VT^1199^) and PER2 (^1205^I**D**VT^1208^). PER1 additionally contains a potential WIN motif (^1109^AA**R**ART^1114^) N-terminal to the CBD, that is not conserved in PER2 (Fig. 1a, Extended Data Fig. 3a). Instead, PER2 contains a WIN motif related sequence (^632^AA**R**PS^636^) between the PAS domains and the CK1δ/ε binding region (Fig. 1a).

**Figure 2:**
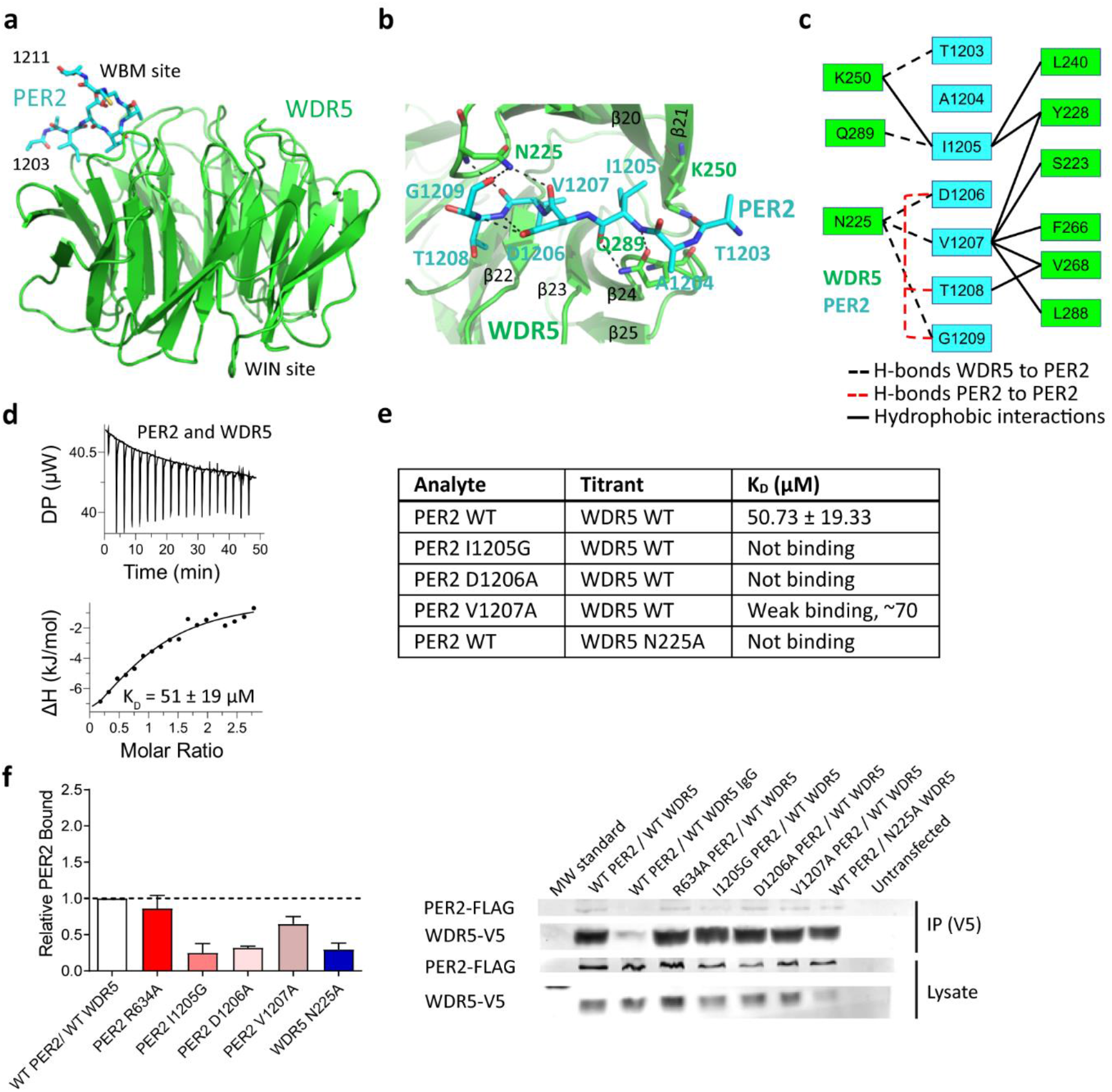
The PER2 WBM motif mediates WDR5 interactions of the purified PER2-CBD protein and of full-length PER2 in HEK293 cells. **a-c** Crystal structure of WDR5 23-334 (green) bound to the PER2 1198-1211 WBM motif peptide (cyan). **a** Overview of the PER2/WDR5 complex. **b** Close-up view of the PER2-WDR5 interaction site. **c** Schematic representation of PER2-WDR5 interactions. Hydrogen bonds were defined by a maximum donor-acceptor distance of 3.1 Å, hydrophobic interactions by a maximum distance of 4.1 Å. **d** ITC experiment of wildtype (WT) PER2 1132-1252 (CBD) and WDR5 23-334. *Upper panel*: raw data, *lower panel*: integrated heat (points) with fit for a one-site binding model (line). Titrant: WDR5. **e** ITC results for interactions of PER2 1132-1252 (WT, I1205G, D1206A, V1207A) with WDR5 23-334 (WT, N225A) as titrant. The K_D_ value for PER2 WT is the means +/- SD from triplicates (Extended Data Table 2, Extended Data Fig. 1, 2a). **f** Co-IP of Flag-tagged full-length PER2 (WT, R634A, I1205G, D1206A, V1207A) with V5-tagged WDR5 (WT, N225A) in HEK293 cells. *Left:* relative PER2 bound to WDR5, normalized to protein expression. Error bars: +/- SEM of 3 replicates*. Right:* Co-IPs with anti-V5 antibody. WDR5-bound Flag-PER2 was quantified by western blot.

**Figure 3:**
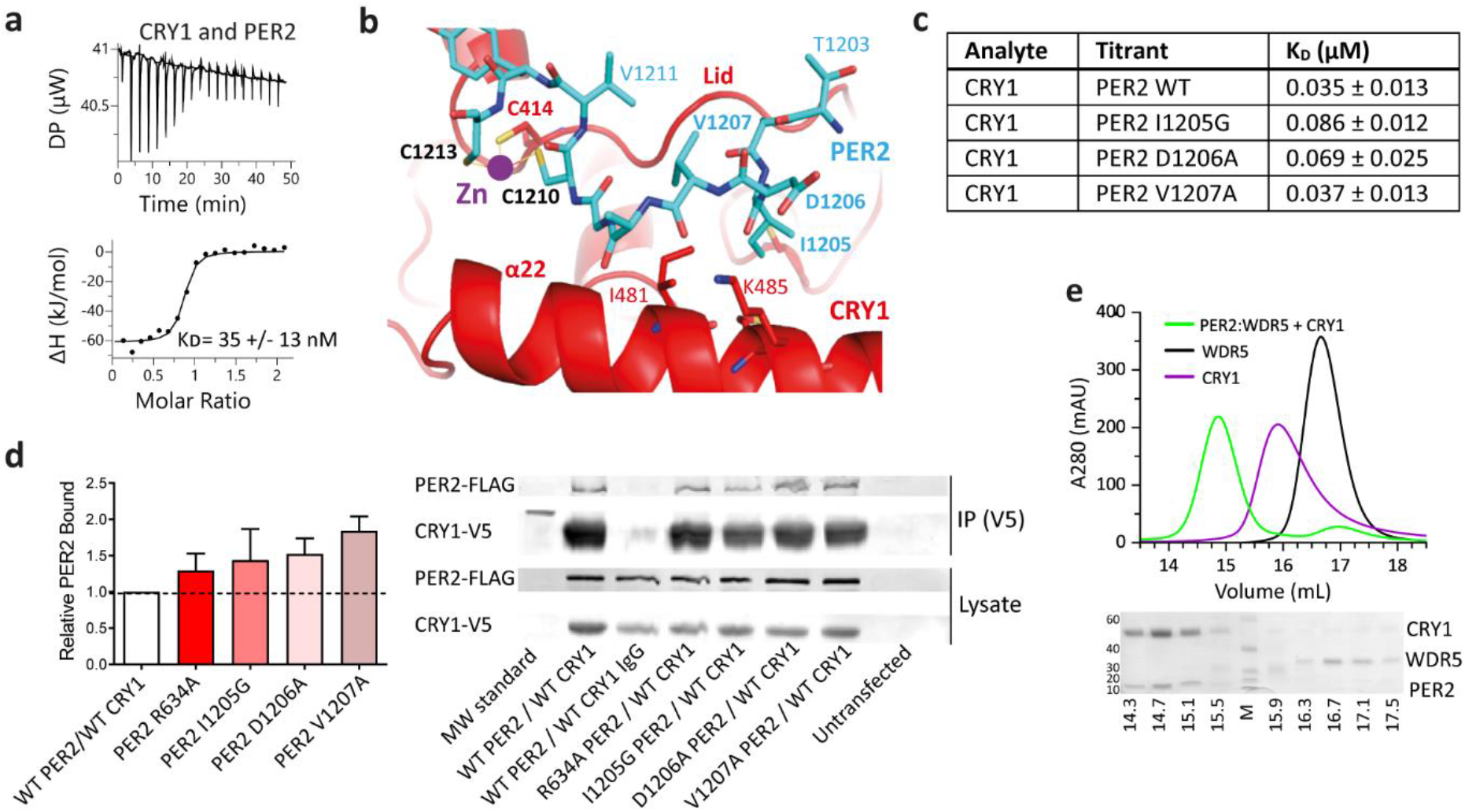
Role of PER2 WBM motif for CRY interactions and competition of WDR5 and CRY1 for PER2 binding. **a** ITC experiment for PER2 1132-1252 WT (CBD) with CRY1 1-496 (PHR). *Upper panel*: raw data, *lower panel*: integrated heat (points) with fit for a one-site binding model (line). Titrant: PER2 **b** Close up view of PER2 WBM motif region (cyan) bound to CRY1-PHR (red) (PDB: 4CT0, ^21^). **c** ITC results (K_D_ values) for binding of PER2 1132-1252 WT, I1205G, D1206A and V1207A (titrant) to CRY1 PHR (mean K_D_ values +/- SD from triplicates) (Extended Data Table 2, Extended Data Fig. 3 b-d). **d** Co-IP results of Flag-tagged PER2 (WT, R634A, I1205G, D1206A, V1207A) with V5-tagged CRY1 in HEK293 cells*. Left:* relative PER2 bound to CRY1 normalized to protein expression. Error bars: +/- SEM of 3 replicates. *Right:* Co-IPs with anti-V5 antibody. CRY1-bound Flag-PER2 was quantified by western blot. **e** SEC analysis (S200 10/300) of WDR5 23-334, CRY1 1-496 and PER2 1132-1252. *Top:* chromatograms of WDR5 (black), CRY1 (magenta) and of a preformed PER2/WDR5 complex (PER2:WDR5) mixed with CRY1 (green). *Bottom:* SDS-PAGE of SEC run with the PER2:WDR5 + CRY1 mixture. No trimeric PER2/WDR5/CRY1 complex was observed, but only a PER2/CRY1 complex eluting at 14.3 to 15.1 mL. WDR5 is displaced by CRY1 and co-elutes with monomeric WDR5 (16.3 to 17.5 mL).

Using analytical size exclusion chromatography (SEC), we showed that purified WDR5 23-334 directly interacts with a C-terminal CRY binding PER2 1132-1252 fragment ^21^ (Fig. 1d). Likewise, a purified C-terminal PER1 1013–1291 fragment including the WDR5 binding WBM- and WIN motifs forms a stable complex with WDR5 in SEC (Fig. 1e). This fragment also includes the PER1 1027-1291 region that interacts with CRY1 in cell-based Co-IP studies ^23^. Hence, we identified WDR5 as a new direct binding partner of PER1 and PER2, targeting the C-terminal PER region that also interacts with CRY.

### PER2 interacts with WDR5 via the WBM motif

To verify that PER2-CBD/WDR5 complex formation is mediated by interactions of the PER2 WBM motif with the WBM site of WDR5, we determined the crystal structure of a complex of WDR5 with a PER2 WBM motif peptide (Y^1198^TGGLPTAI**D**VTGCV^1211^) (Fig. 2 a-c, Extended Data Table 1). Residues T1203 to V1211 of the PER2 peptide are well resolved in the electron density (Extended Data Fig. 1a). Our 3D-structure revealed that the PER2 WBM motif ^1205^I**D**VTG^1209^ indeed binds to the WBM motif binding pocket of WDR5, inserting Ile1205 and Val1207 into its two hydrophobic sub-pockets (Fig. 2b). The side chain of the highly conserved aspartate (Asp1206 in PER2) forms intramolecular hydrogen bonds to the PER2 backbone amino groups of Thr1208 and Gly1209. Additionally, Asn225, Gln289 and Lys250 of WDR5, which are located in two WDR5 loops flanking the WBM motif binding pocket, form hydrogen bonds to the PER2 backbone carbonyl oxygens of Gly1209, Val1207, Asp1206 (Asn225) and Thr1203 (Lys250) as well as to the backbone carbonyl and amino groups of Ile1205 (Gln289) (Fig. 2 b,c). Overall, the WBM motif of PER2 binds WDR5 in a similar location and orientation as the WBM motif of RbBP5 ^29^ (Extended Data Fig. 1b).

To show that the interaction between the PER2 WBM motif and the WBM site of WDR5 also occurs in solution and in the context of the PER2-CBD, we mutated three WDR5 interacting amino acids of PER2 (Ile1205, Asp1206, Val1207) as well as Asn225 of WDR5 to alanine (D1206A, V1207A, N225A) or glycine (I1205G) and analyzed the interactions between PER2 1232-1252 (CBD) and WDR5 23-334 (wildtype and mutants) by SEC and Isothermal titration calorimetry (ITC) (Fig. 2 d,e, Extended Data Fig. 1c-g, 2a, Extended Data Table 2). Although the PER2-CBD and WDR5 form a stable complex in SEC (Fig. 1d), their interaction affinity is rather low with a K_D_ value of about 50 µM (Fig. 2d). The D1206A mutation completely disrupted the PER2/WDR5 complex in SEC and ITC, suggesting that the intramolecular interactions of Asp1206 are important, likely by stabilizing the PER2 conformation required for WDR5 binding. The I1205G mutation also completely disrupted the PER2-WDR5 interaction in SEC and ITC, while the V1207A mutation led to a partial dissociation of the complex in SEC and a very low WDR5 affinity (K_D_ about 70 µM) (Fig. 2e, Extended Data Fig. 1c-g). Furthermore, the WDR5 N225A mutation eliminates PER2 binding in ITC (Fig. 2e, Extended Data Fig. 2a), confirming that PER2 binds to the WBM site of WDR5. We also crystallized the WDR5 N225A mutant protein and could see that the overall structure of WDR5 was not affected beyond the region around the mutation site (Extended Data Fig. 2b, Extended Data Table 1).

To validate the PER2-WDR5 interface within full-length proteins and in a cellular context, we performed Co-Immunoprecipitation (Co-IP) experiments of overexpressed full-length V5-tagged WDR5 (wildtype, N225A) and Flag-tagged PER2 (WT, R634A, I1205G, D1206A, V1207A) proteins in HEK293 cells. Co-IPs were carried out with an anti-V5 antibody and bound PER2 was analyzed by western blot with an anti-Flag antibody (Fig. 2f). We found, that the PER2 I1205G and D1206A mutations as well as the WDR5 N225A mutation led to a significantly reduced PER2 binding compared to the wild-type PER2-WDR5 interaction (about a quarter of PER2 binds to WDR5 compared to wildtype proteins), while about half of the PER2 V1207A mutant protein binds to WDR5 compared to wildtype. These results are in very good qualitative agreement with our SEC and ITC experiments and show that full-length PER2 also interacts with the WDR5 WBM site via the WBM motif in cells. Notably, PER2 R634A with a mutation of the conserved arginine in a potential WIN motif (^633^A**R**PS^636^) (Fig. 1a) showed a wild-type like binding to WDR5 in our Co-IP studies (Fig. 2f). Hence, the only potential WIN motif of PER2 is not required for the WDR5-PER2 interaction. Collectively our data show that the ^1205^IDVT^1208^ WBM motif of PER2 mediates interactions with the WBM site of WDR5 in cells (full-length PER2) and within the purified PER2-CBD protein.

### The PER2 WBM motif binds CRY1 near the Zinc interface, but WBM mutations do not disrupt CRY1-PER2 interactions

Interestingly, the PER2 WBM motif is located within the CRY1-PER2 interface, between helix α22 and the “Lid” loop of the CRY1 photolyase homology region (PHR) and next to a zinc ion that is jointly coordinated by Cys1210/Cys1213 of PER2 and Cys414/His473 of CRY1 (Fig. 3b, Extended Data Fig. 3a) ^21^. To elucidate the role of the WBM motif in PER2-CRY1 interactions, we performed ITC experiments with PER2 1132-1252 (WT, I1205G, D1206A, V1207A) and CRY1 1-496, which includes the PHR region of CRY1, shown to be sufficient to bind PER2 ^21^ (Fig. 3 a-c). All PER2 proteins bound to CRY1 with a high nM affinity with K_D_ values of 35 nM (wildtype), 86 nM (I1205G), 69 nM (D1206A) and 37 nM (V1207A) (Fig. 3 a,c; Extended Data Fig. 3 b-d). Hence, the WBM motif mutations, while weakening or disrupting WDR5-PER2 interactions, do not impact the CRY1 binding affinity much, despite the striking location of the WBM motif near the zinc interface. This is likely due to the large CRY1-PER2 interface area ^21^ where the PER2 WBM motif only provides a minor contribution, as well as the much higher affinity of PER2 to CRY1 (nM) compared to WDR5 (µM).

We also performed Co-IP experiments for full-length PER2 (WT, R634A, I1205G, D1206A, V1207A) and CRY1. In contrast to their weakening effect on PER2-WDR5 interactions (Fig. 2f), WBM motif mutations did not reduce PER2 binding to CRY1 in HEK293 cells (Fig. 3d). This result is consistent with our ITC studies and shows, that – despite its location in the CRY1-PER2 interface - an intact WBM motif is not essential for CRY1-PER2 interactions in cells, likely due to the higher affinity and additional binding sites in the CRY1/PER2 complex. Furthermore, the R634A mutation in the potential PER2 WIN motif did not impact CRY1 binding, as expected based on its location remote from the C-terminal CRY binding domain (Fig. 3d).

### CRY1 and WDR5 compete for PER2 binding

To assess if WDR5 and CRY1 compete for PER2 binding due to their overlapping binding sites in the C-terminal PER2 CBD, we performed SEC experiments with a pre-purified PER2-CBD/WDR5 complex incubated with equimolar CRY1 1-496 (PHR) (Fig. 3e) as well as with an equimolar mixture of CRY1-PHR, PER2-CBD and WDR5 proteins (data not shown). A trimeric PER2/WDR5/CRY1 complex was not observed in any of the experiments, but only a dimeric PER2/CRY1 complex. Therefore, WDR5 and CRY1 indeed compete for binding to PER2. Moreover, CRY1 sequesters PER2 from a preformed PER2/WDR5 complex, consistent with its much higher affinity for PER2.

### PER1 has a higher WDR5 affinity than PER2 and forms a tight complex with CRY1

Using SEC and ITC we showed that the C-terminal PER1 1013-1291 fragment directly interacts with WDR5 23-334 (Fig. 1e; Fig. 4a,c,d) and with CRY1 1-496 (PHR) (Fig. 4 b,c,e). Interestingly, PER1 binds to WDR5 with a K_D_ value of 3.1 µM (Fig. 4a,c), which corresponds to a ∼15 times higher affinity than PER2 (∼50 µM) (Fig. 2d,e). Furthermore, PER1 1013-1291 binds to CRY1 with a K_D_ value of 16.4 nM (Fig. 4b,c), i.e. with a similar affinity as PER2 (Fig. 3a,c). The comparably high nM range affinities of PER1 and PER2 for CRY1 are in sharp contrast to their overall much lower (µM range) and unequal affinities for WDR5.

**Figure 4:**
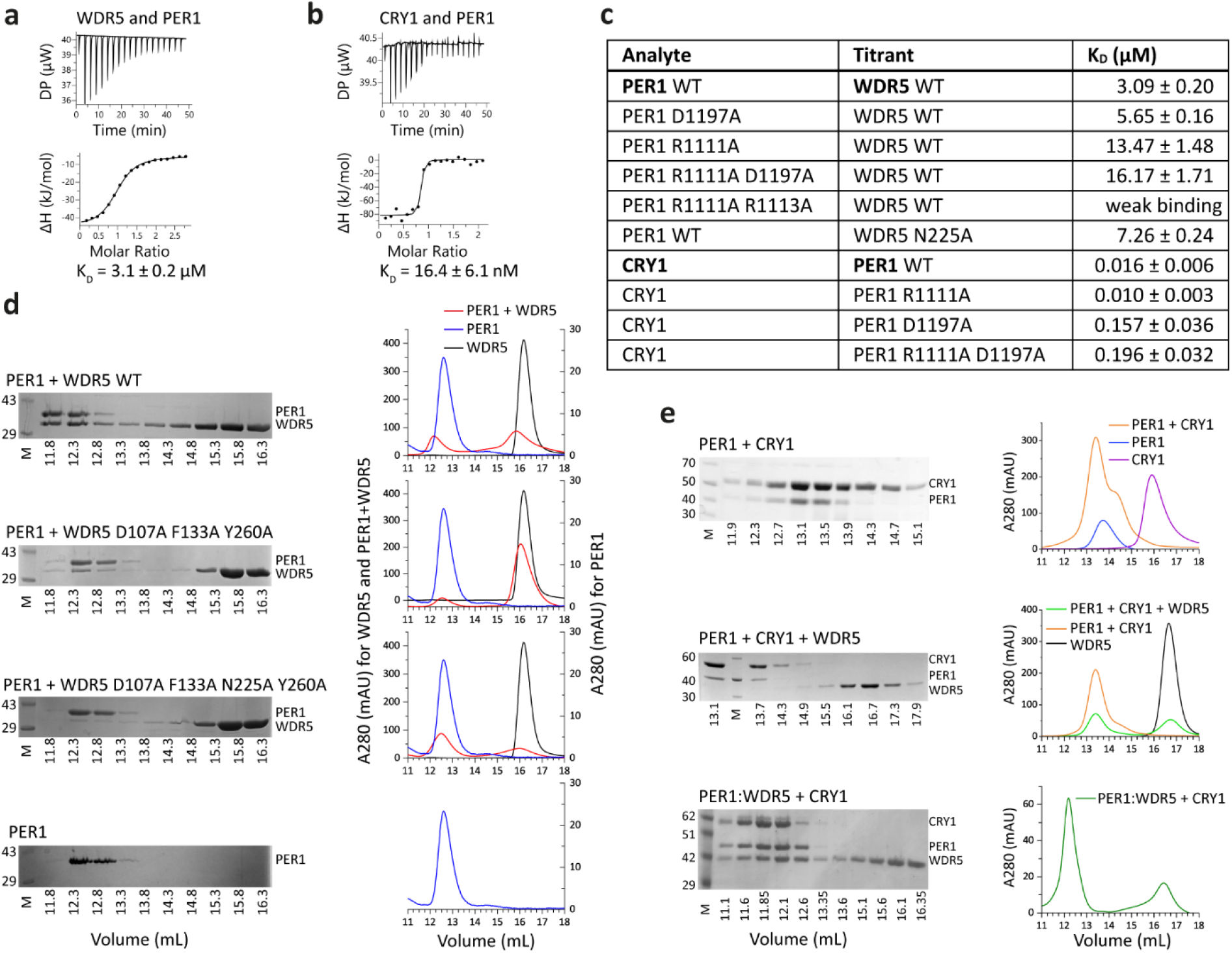
PER1 predominantly binds to the WDR5 WIN site and can form a trimeric PER1/WDR5/CRY1 complex. **a,b** ITC analyses of the interaction of PER1 1013-1291 with **a** WDR5 23-334 (titrant WDR5) and with **b** CRY1 1-496 (PHR) (titrant PER1). Data were fit with a one-site binding model (lower panel). **c** ITC results (K_D_) for interactions of PER1 1013-1291 (WT, R1111A, D1197A, R1111A D1197A, R1111A R1113A) with WDR5 23-334 (WT, N225A) or CRY1 1-496 (mean K_D_ values +/- SD from triplicates) (Extended Data Table 2, Extended Data Fig. 4 a-d and 5 a-c). **d** SEC (S200 10/300) analyses of the interaction of PER1 1013-1291 with WDR5 23-334 WT (*top),* the WIN site triple mutant WDR5 D107A F133A Y260A and the WIN/WBM site mutant WDR5 D107A F133A Y260A N225A. The SEC run of PER1 alone is shown for comparison *(bottom). Left:* SDS-PAGE analyses of peak SEC fractions, *right:* SEC chromatograms of the PER1/WDR5 complex (red), PER1 alone (blue) and WDR5 alone (black). **e** SEC (S200 10/300) analyses of the PER1 1013-1291 interaction with CRY1 1-496 and WDR5 23-334. *Top, left*: SDS-PAGE of PER1/CRY1 complex SEC, *right:* chromatograms of PER1 (blue), CRY1 (magenta) and PER1/CRY1 complex (orange). *Middle, left:* SDS-PAGE of SEC with equimolar mixture of PER1, CRY1 and WDR5, *right:* chromatograms of WDR5 (black), PER1/CRY1 complex (orange) and CRY1+PER1+WDR5 mixture (green). Only a dimeric PER1/CRY1 complex is observed, WDR5 is displaced. *Bottom:* SDS-PAGE *(left)* and chromatogram *(right)* of a preformed PER1/WDR5 complex (PER1:WDR5) incubated with equimolar CRY1 (green). A trimeric PER1/WDR5/CRY1 complex is formed.

### The WIN site is more important for PER1-WDR5 interactions than the WBM site

To find out, if the potential WBM and WIN motifs of PER1 (Fig. 1a,b; Extended Data Fig. 3a) are involved in WDR5 binding, we mutated the conserved Arg1111 (WIN) and Asp1197 (WBM) to Ala and determined the WDR5 binding affinities by ITC. The single mutations increased the K_D_ values to 13.5 µM (R1111A) and 5.7 µM (D1197A) respectively, the R1111A D1197A double mutant to 16 µM (Fig. 4c, Extended Data Fig. 4a-c). Given the rather mild effects of these mutations, we hypothesized that the second arginine, Arg1113, present in the potential WIN motif of PER1 could also play a role in WDR5 binding, akin to the KANSL1/WDR5 WIN site interaction in the non-specific lethal (NSL) complex (PDB 4CY2, ^36^). In the KANSL1/WDR5 complex, the first highly conserved arginine of the KANSL1 WIN motif (corresponding to Arg1111 in PER1) is inserted in the WIN motif binding pocket of WDR5 nearby Phe133, whereas the second arginine (Arg1113 in PER1) forms a salt bridge to Asp107_WDR5_ at the surface of the WDR5 WIN site (Extended Data Fig. 4e). Indeed, the R1111A R1113A double mutation drastically reduces the WDR5 affinity of the PER1 CBD (K_D_ > 100 µM) (Fig. 4c, Extended Data Fig. 4d), suggesting that Arg1113 is important for PER1 binding to the WDR5 WIN site. To confirm that PER1 binds to the WIN site of WDR5, we generated – based on the KANSL1/WDR5 complex crystal structure (Extended Data Fig. 4e) - a WDR5 D107A F133A Y260A triple mutant protein designed to disrupt interactions with PER1 residues R1111 (F133A), R1113 (D107A) and T1114 (Y260A). The triple mutant mostly disrupted the WDR5/PER1 complex in SEC (Fig. 4d), confirming the importance of the WDR5 WIN site for PER1 binding. Moreover, the simultaneous disruption of the WIN- and WBM sites of WDR5 by combining the D107A F133A Y260A triple mutant (WIN site) with the N225A mutation (WBM site) leads to a complete dissociation of the PER1/WDR5 complex in SEC (Fig. 4d), suggesting that the WBM site is also involved in PER1 binding. The moderate effects of the D1197A mutation in the PER1 WBM motif (K_D_ = 5.7 µM) and of the single WDR5 N225A mutation (K_D_ ∼ 7 µM) on the PER1-WDR5 interaction affinity (Fig. 4c, Extended Data Fig. 2c) as well as the very low WDR5 affinity of the PER1 R1111A R1113A WIN site mutant (K_D_ > 100 µM) however indicate a minor importance of the WBM site for PER1-WDR5 complex formation. This contrasts with PER2 and RbBP5, which bind WDR5 exclusively via their WBM motif, leading to a much more drastic effect of the WDR5 N225A mutation by disrupting PER2-WDR5 interactions (Fig. 2e, Extended Data Fig. 2a) and lowering the RbBP5-WDR5 affinity from a K_D_ value of 0.62 µM (wildtype WDR5) to 71 µM (WDR5 N225A) (Extended Data Fig. 2 d,e). Together, our data show that PER1 binds to both, the WIN- and the WBM motif binding pockets of WDR5, but the WIN site interaction is more important for complex formation than the WBM site.

### The PER1 WBM motif is involved in CRY1 binding

We also tested the impact of the PER1 R1111A WIN - and D1197A WBM motif mutations on CRY1 binding. The PER1 1013-1291 R1111A protein had a similar affinity for CRY1 1-496 (PHR) as wildtype PER1, whereas the D1197A mutation lead to a ten-fold higher K_D_ value of 157.3 nM (Fig. 4c, Extended Data Fig. 5a-c). The R1111A D1197A double mutation affecting the WIN- and WBM motif resulted in a K_D_ value of 196 nM, comparable to the D1197A single mutation. Hence, the PER1 WBM motif is involved in CRY1 binding, while the WIN motif region is not required for PER1-CRY1 interactions. This result is consistent with the predicted CRY1 binding site of PER1 (based on the CRY1/PER2 crystal structure ^21^), which includes the WBM-but not the WIN motif (Extended Data Fig. 3a). Notably, the significant impact of the D1197A mutation on the PER1-CRY1 interaction (10-fold reduced affinity) contrasts with the minor effect of the analogous D1206A_PER2_ mutation on PER2-CRY1 interactions (Fig. 3c, Extended Data Fig. 3c) suggesting a more important role of this interface in the PER1/CRY1 complex.

**Figure 5:**
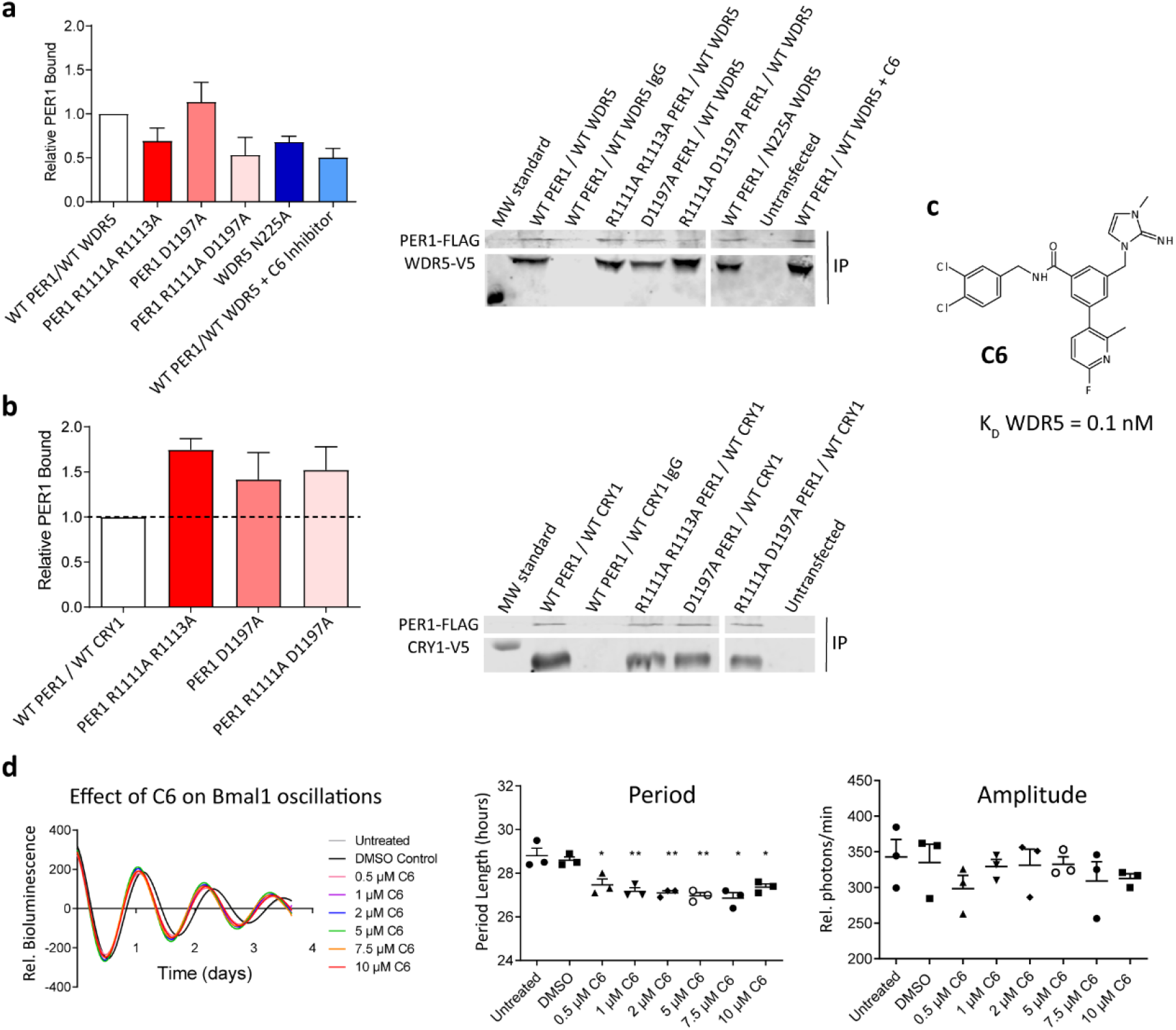
Effect of WIN- and WBM interface mutations and C6 WIN site inhibitor on PER1-WDR5- and PER1-CRY1 interactions in HEK293 cells and on circadian oscillations in U2OS cells **a** Co-IP of Flag-tagged full-length PER1 (WT, R1111A R1113A, D1197A, R1111A D1197A) with V5-tagged WDR5 (WT, N225A) in HEK293 cells. The C6 compound was added to wildtype PER1 and WDR5 at a 5 µM concentration. *Left: R*elative PER1 bound to WDR5 normalized to protein expression. *Right:* Co-IPs with anti-V5 antibody. WDR5-bound Flag-PER1 was quantified by western blot. **b** Co-IP of Flag-tagged PER1 (WT, R1111A R1113A, D1197A, R1111A D1197A) with V5-tagged CRY1 in HEK293 cells. *Left:* Relative PER1 bound to CRY1 normalized to protein expression*. Right:* Co-IPs with anti-V5 antibody. CRY1-bound Flag-PER1 was quantified by western blot. Error bars in a,b: +/- SEM of 3 replicates. **c** Chemical structure of the C6 WIN site inhibitor ^33^. **d** C6 shortens the period of circadian Bmal1-Luc oscillations in U2OS cells by 2 to 3 h at all tested concentrations but does not significantly change the oscillation amplitude. *Left:* average relative bioluminescence n=3, *middle:* Period length, * p<0.05, ** p<0.01, two-tailed unpaired t test, *right:* relative amplitude (arbitrary units), no significant differences, two-tailed unpaired t test.

### Trimeric PER1/WDR5/CRY complexes can be sequentially assembled

As WDR5 predominantly binds to the WIN motif of PER1, which is not required for CRY interactions, we hypothesized that a trimeric PER1/WDR5/CRY complex could be formed. Using equimolar mixtures of purified WDR5 23-334, PER1 1013-1291 and CRY1 1-496 proteins, we did not observe a trimeric WDR5/PER1/CRY1 complex in SEC, but only a dimeric PER1/CRY1 complex with displacement of monomeric WDR5 (Fig. 4e, middle). However, we did observe a trimeric PER1/WDR5/CRY1 complex, when CRY1 was added to a preformed PER1/WDR5 complex (Fig. 4e, bottom). Moreover, we observed the same behavior for the CRY2-PHR (1-512): a trimeric PER1/WDR5/CRY2 complex was only formed when CRY2 was added to a preformed PER1/WDR5 complex (Extended Data Fig. 5d). Given that CRY has a higher PER1 affinity than WDR5 (Fig. 4a-c), the PER1 WIN motif is located directly N-terminal to the PER CBD (Extended Data Fig. 3a) and the CBD undergoes an induced folding upon CRY binding (Schmalen et al, 2014), it is conceivable that CRY/PER1 complex formation traps the PER1 WIN motif (preventing its accessibility for WDR5), unless the WIN motif is priorly associated with WDR5 in a preassembled PER1/WDR5 complex. The formation of trimeric PER1/WDR5/CRY1 complexes from preassembled PER1/WDR5 complexes may in turn be supported by the significant contribution of the PER1 WBM motif to PER1-CRY complex formation (as evidenced by the D1197A mutation, Fig. 4c), which could facilitate a local displacement of the PER1 WBM motif from WDR5 by CRY, without disrupting preexisting WDR5 interactions with the PER1 WIN motif. Together, our data show that the unique features and locations of the WIN- and WBM motifs as well as the different WDR5 interaction interfaces of PER1 and PER2 translate into distinct interaction interplays with CRY: while PER2 can only form dimeric complexes with either WDR5 or CRY, PER1 is able to sequentially assemble into trimeric PER1/WDR5/CRY complexes.

### WIN motif mutations and a WDR5 WIN-site inhibitor weaken PER1-WDR5 interactions in HEK293 cells

To validate the PER1-WDR5 interface within full-length proteins and in a cellular context, we performed Co-IP experiments of overexpressed V5-tagged WDR5 (WT, N225A) and Flag-tagged PER1 (WT, D1197A, R1111A D1197A, R1111A R1113A) proteins in HEK293 cells. Co-IPs were carried out with an anti-V5 antibody and bound PER1 was analyzed by western blot (Fig. 5a). We found, that the single PER1 D1197A WBM motif mutation did not affect PER1-WDR5 interactions in cells. In contrast, the PER1 R1111A R1113A (WIN) and R1111A D1197A (WIN/WBM) double mutations significantly weakened the interaction. These findings show that the PER1 WIN motif is more important than the WBM motif for the interaction of full-length PER1 with WDR5 in cells. Moreover, they illustrate the different binding modes of PER1 and PER2 to WDR5 in cells (Fig. 2f, Fig. 5a).

As PER1 predominantly targets the WIN site of WDR5, we also tested the effect of a WIN site inhibitor (C6, Fig. 5c), that binds to the WDR5 WIN site with a ∼100 pM affinity and is considered for treatment of MLL1- and Myc-related cancers ^33^, on the PER1-WDR5 interaction by Co-IP. Interestingly, C6 significantly weakens the PER1-WDR5 interactions in HEK293 cells (Fig. 5a), further confirming the importance of the WIN site for WDR5/PER1 complex formation.

Notably, the WDR5 N225A WBM site mutation also reduces PER1 binding in our Co-IP experiments. This contrasts with the moderate effect of the N225A mutation on the purified WDR5/PER1-CBD complex (Fig. 4c, Extended Data Fig. 2c) and with the minor effect of the PER1 D1197A mutation on WDR5-PER1 interactions *in vitro* (Fig. 4c, Extended Data Fig. 4a) and *in vivo* (Fig. 5a). Given the known scaffold function of WDR5 in other cellular multi-protein complexes ^24^, we propose, that in cells the WBM binding pocket of WDR5 could mediate interactions with another (yet unknown) protein, that stabilizes the PER1-WDR5 interaction within a larger complex.

None of the tested WIN- and WBM motif mutations weakened the PER1-CRY1 interactions in HEK293 cells (Fig. 5b). Although the D1197A mutation reduces the CRY1-PER1 CBD interaction affinity from 16 nM to 157 nM (Fig. 4c, Extended Data Fig. 5b,c), the interaction remains at a wild-type like level in the Co-IP experiment, likely stabilized by other binding sites that are not affected by the D1197A mutation. It is however also possible, that the amounts of CRY1 and PER1 proteins in the cell were too high to observe affinity differences in the nanomolar range in the western blot analyses of the Co-IP experiment.

### The WDR5 WIN site inhibitor shortens the circadian period in U2OS cells

To find out if C6 impacts circadian clock function, we also tested the effect of the C6 compound on circadian Bmal1-Luc oscillations in U2OS cells. Interestingly, C6 significantly shortened the period of circadian oscillations by 2 to 3 h at all tested compound concentrations from 0.5 µM to 10 µM C6 without affecting the oscillation amplitude. This effect was not observed for the DMSO control (Fig. 5d). Hence, WDR5 and its WIN site interactions with PER1 or with other WIN site ligands are important for circadian clock function. Apart from PER1, MLL1 constitutes another WDR5 WIN site ligand within the mammalian circadian clock, that could in principle be targeted by the C6 compound. However, the short period caused by the C6 compound treatment is more consistent with the short period phenotype resulting from PER1 disruption ^14–16^ than with MLL1 inactivation ^26^. Furthermore, the C6 compound does not disrupt WDR5-MLL1 interactions in HEK293 cells and is thought to affect cells independent of MLL1 ^33,37^. While we cannot exclude that C6 targets an unknown WDR5 WIN site ligand with a clock-related function, our study identifies PER1 as a plausible target mediating the observed period shortening by the C6 compound, as i) PER1 binds to the WDR5 WIN site and C6 disrupts the PER1-WDR5 interaction in HEK293 cells (this study), and ii) PER1 disruption causes a short period phenotype in mice and cells ^14–16^. In conclusion, by uncovering a hitherto unknown and unexpected effect of C6 on circadian clock function, our study broadens the scope of potential cellular targets of this class of WIN site inhibitors and sets the stage to further explore their mechanism of action within the circadian clock.

### A trimeric PER1/WDR5/PER2 complex is not observed *in vitro*

Our observation that the WDR5 N225A mutation weakens PER1-WDR5 interactions in cells, but only moderately reduces the WDR5 affinity of the purified PER1-CBD fragment, prompted us to speculate that in cells another protein binds to the WDR5 WBM site together with WIN site bound PER1 to form a trimeric complex with enhanced stability. Thereby, the N225A mutation could weaken PER1-WDR5 interactions by displacing this additional WBM site ligand. As PER2 only binds to the WBM site of WDR5, we hypothesized that a trimeric PER1/WDR5/PER2 complex could be formed with PER1 binding to the WIN site and PER2 to the WBM site, potentially enhancing the stability and specificity of the complex by additional PER1-PER2 interactions, e.g. via the PAS domains ^18,19,38^. To test this hypothesis, we conducted SEC experiments with our purified PER1 1013-1291, PER2 1132-1252 (CBD) and WDR5 23-334 proteins. In the SEC, we did not observe a trimeric PER1/WDR5/PER2 complex, but rather found WDR5 binding to either PER1 or PER2 (Fig. 6a). The dimeric PER1/WDR5 and PER2/WDR5 complexes eluted in two well separated peaks at their expected elution volumes of about 13.5 mL (PER1/WDR5) and 15.5 mL (PER2/WDR5) (compare Fig. 1e, Extended Data Fig. 1d, Fig. 6b) and earlier than free WDR5 (∼ 16.5 mL). The absence of PER1/WDR5/PER2 trimers could be due to steric hindrances but also be related to the low WDR5 binding affinity of the PER2-CBD, which may prevent it from displacing PER1 from its minor interaction site at the WBM binding pocket. Certainly, our observation argues against a stabilization of the PER1/WDR5 complex by PER2, at least in the context of the C-terminal WDR5- and CRY binding PER regions.

**Figure 6:**
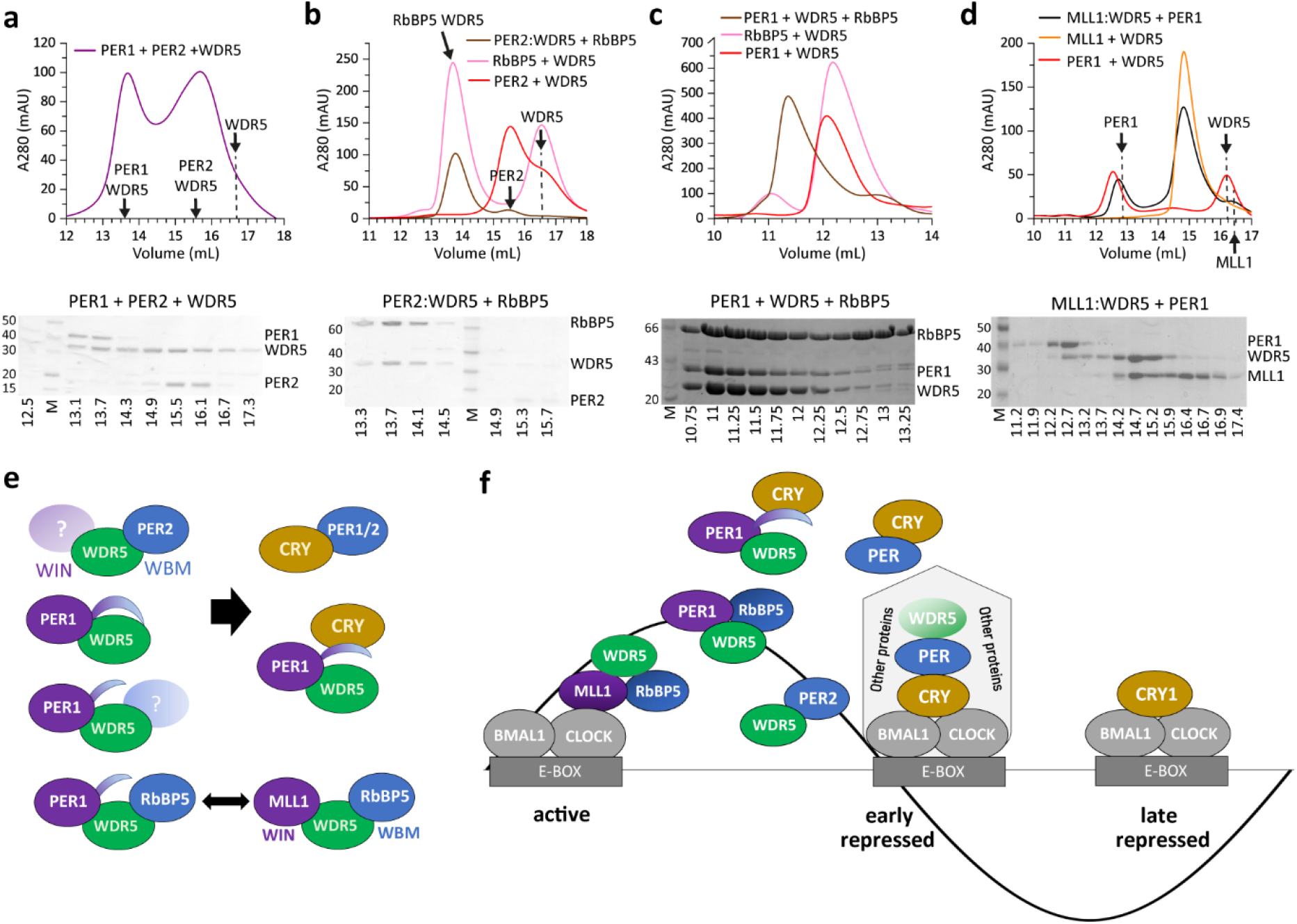
Interaction interplay between PER, WDR5, RbBP5, MLL1 and CRY. **a** Analytical SEC (S200 10/300) of PER1 1013-1291, PER2 1132-1252 and WDR5 23-334. Chromatogram *(top)* and SDS-PAGE *(bottom)* of the SEC run with the PER1+WDR5+PER2 mixture. No trimeric PER1/WDR5/PER2 complex was observed, but only dimeric PER1/WDR5 and PER2/WDR5 complexes. Arrows indicate elution volumes of individual PER1/WDR5- and PER2/WDR5 complexes and of WDR5. **b** Analytical SEC (S200 10/300) of PER2 1132-1252, RbBP5 and WDR5 23-334. *Top:* chromatograms of PER2/WDR5 (red)- and RbBP5/WDR5 (magenta) complexes (excess WDR5) and of a competition experiment of a preformed PER2/WDR5 complex (PER2:WDR5) with added equimolar RbBP5 (brown). *Bottom:* SDS-PAGE of the competition experiment (SEC peak fractions). No trimeric PER2/WDR5/RbBP5 complex was observed, but only the dimeric WDR5/RbBP5 complex. PER2 is displaced by RbBP5. **c** Analytical SEC (S200 10/300) of PER1 1013-1291, RbBP5 and WDR5 23-334. *Top:* chromatograms of PER1/WDR5 (red)-, RbBP5/WDR5 (magenta) and PER1/WDR5/RbBP5 (brown) complexes. *Bottom:* SDS-PAGE of the trimeric PER1/WDR5/RbBP5 complex (SEC peak fractions), that eluted significantly earlier than the dimeric complexes and the individual PER1 and WDR5 proteins (Fig. 4d). Note that a different SEC S200 10/300 column was used in Fig. 4d and Fig. 6c,d than in all other experiments stating S200 10/300. **d** Analytical SEC (S200 10/300) of WDR5 23-334, PER1 1013-1291 and hMLL1 3754-3963 (MLL1-WIN-SET). *Top:* chromatograms of PER1/WDR5 (red)- and MLL1/WDR5 (orange) complexes and of the competition experiment of a preformed MLL1/WDR5 complex (MLL1:WDR5) with added equimolar PER1 (black). *Bottom:* SDS-PAGE of the competition experiment. Newly formed PER1/WDR5 complex elutes at ∼ 12.5 mL, remaining MLL1/WDR5 complex at ∼ 15 mL. PER1 partially displaces MLL1 (∼ 16.5 mL) from the preformed MLL1/WDR5 complex. **e** Schematic of PER-WDR5 interactions and their interplay with CRY, RbBP5 and MLL1. PER2 only binds to the WBM site of WDR5 and another protein could bind to the WDR5 WIN site. PER1 binds to both WDR5 binding sites via its WIN- and WBM motifs, but the WIN site interaction is more important. Alternatively, PER1 can only bind to the WIN site, while another protein binds to the WBM site, for example RbBP5. Thereby PER1 could replace MLL1 on the WIN site. PER2 binds to either WDR5 or CRY, whereas PER1 can form a trimeric PER1/WDR5/CRY complex. **f** Active-, early repressed- and late repressed states of BMAL1/CLOCK in the mammalian circadian oscillator. WDR5 is an MLL1 complex component associated with active BMAL1/CLOCK, but was also identified in a repressive PER/CRY containing complex. PER1/WDR5/RbBP5- and PER1/WDR5/CRY complexes may be formed at the transition from the active-to the early repressed state. PER2/WDR5 complexes occur in a CRY-independent manner.

### PER1 displaces MLL1 from WDR5 and forms a trimeric complex with WDR5 and the MLL1 complex component RbBP5

In the MLL1 complex, which co-activates BMAL1/CLOCK, RbBP5 binds to the WBM site of WDR5, while the WIN binding pocket is occupied by MLL1 ^28–32^. RbBP5 binds WDR5 with a K_D_ of 0.6 µM, i.e. about 80 times stronger than PER2 (Fig. 2d, 4a; Extended Data Fig. 2d). Consistent with its higher WDR5 affinity and binding competition with PER2 for the WBM site, RbBP5 displaces the PER2-CBD from a preformed PER2/WDR5 complex to form a dimeric RbBP5/WDR5 complex, and a trimeric PER2/WDR5/RbBP5 complex cannot be formed (Fig. 6b). In contrast, we could observe a trimeric complex between WDR5, RbBP5 and the PER1-CBD in SEC, suggesting that in presence of RbBP5, PER1 can bind to WDR5 independent of its WBM motif via the WIN pocket of WDR5 (Fig. 6c). Furthermore, the PER1-CBD partially displaces an MLL1-WIN-SET fragment (MLL1 3754-3963, comprising the WDR5 binding WIN motif and the catalytic SET domain of MLL1) from a preformed MLL1/WDR5 complex due to its competition with the WDR5 WIN site, to form a dimeric PER1/WDR5 complex (Fig. 6d, Extended Data Fig. 6). This finding suggests that PER1 (but not PER2) has the potential to affect the integrity of the MLL1 complex based on its ability to displace MLL1 from the WIN site and to interact with WDR5 in presence of RbBP5.

## Discussion

Our studies revealed a new role of WDR5 as a direct PER interaction partner in the mammalian circadian clock. We uncovered critical differences between the PER1- and PER2 interactions with WDR5, that translate into different molecular interactions at the transition from active- to repressed states of BMAL1/CLOCK. Importantly, we identified two WDR5 binding sites in PER1, that effectively act as a molecular chokepoint allowing sequential complex formations during the transition from circadian gene activation to repression (Fig. 6 e,f).

Within the mammalian circadian clock, WDR5 was identified as part of the BMAL1/CLOCK co-activating MLL1 complex, that promotes open chromatin via histone H3K4 trimethylation, as well as in a repressive PER complex, which is thought to induce a closed chromatin state by histone H3K9 methylation (involving the SUV39H1 methyltransferase) and histone deacetylation ^7,9,10,13,25,26^. Consistent with this dual role, depletion of WDR5 abolishes circadian rhythms of both, methylation of histone H3K4 (activation) and histone H3K9 (repression) ^9^, placing WDR5 in a key role at the interface between circadian gene activation and repression. Here, we analyzed the molecular interactions of WDR5 with PER1 and PER2, which are likely to be involved in the transition towards the repressive state, as PER1 and PER2 proteins accumulate at circadian promotors within this time window ^7–9^. Interestingly, we discovered striking differences between PER1 and PER2 regarding their interaction sites and affinities for WDR5. We showed that PER2 has one WDR5 binding site (the WBM motif), which binds to the WBM pocket of WDR5. Intriguingly, this WBM motif is located within the C-terminal CRY binding domain (CBD) of PER2 ^21^, leading to binding competition between WDR5 and CRY (Fig. 2,3). While this WBM motif is also present in PER1, PER1-WDR5 interactions are predominantly stabilized via a second WDR5 binding site, the WIN motif, which is located outside of the PER1 CBD and binds to the WIN site of WDR5. This WIN site interaction gives PER1 a fifteen-fold higher affinity for WDR5 compared to PER2 and enables it to form trimeric PER1/WDR5/CRY- and PER1/WDR5/RbBP5 complexes (Fig. 4, 5a,b, 6c, Extended Data Fig. 5d). Hence, the unique features and locations of its WIN- and WBM motifs allow PER1 to create two “molecular gates”: the first permitting a trimeric PER1/WDR5/RBP5 early repression complex that would succeed the MLL1/WDR5/RBP5 activation complex at circadian promotors, and the second gate permitting PER1/WDR5/CRY complex formation to transition towards CRY/PER containing repressive complexes (Fig. 6 e,f).

The first gate relies on the ability of PER1 to form a trimeric PER1/WDR5/RbBP5 complex, where PER1 replaces MLL1 on the WDR5 WIN site (Fig. 6c). MLL1 is thought to be recruited to circadian promotors via tandem BMAL1/CLOCK complexes stabilized by the exon19 region of CLOCK ^26,39,40^, suggesting local MLL1 concentrations to be sub-stoichiometric to BMAL1/CLOCK. It is therefore conceivable, that PER1 can – following its circadian accumulation in the nucleus - exceed the local MLL1 concentration on circadian promotors. We therefore propose that the displacement of the catalytic MLL1 subunit by circadianly accumulating PER1 proteins could provide a molecular mechanism to terminate MLL1-dependent histone H3 tri-methylation and transcriptional activation of circadian genes in a timely manner (when the local PER1 concentration is high enough to displace MLL1) to allow the progression towards a repressive chromatin state.

In contrast to MLL1 and WDR5, which do not cycle over the day ^9,26^, RbBP5 mRNA levels are reported to be clock-controlled and to cycle in antiphase to PER and CRY ^41^. Hence, RbBP5 protein levels are likely to decline, leaving binary PER1/WDR5 complexes behind, before the CRY/PER containing early repressive complex assembles. Preformed PER1/WDR5 complexes can then associate with accumulating CRY1/2 proteins to form trimeric PER1/WDR5/CRY complexes (Fig. 4e, Extended Data Fig. 5d), where PER1 binds to WDR5 via the WIN motif and to CRY via the CRY-binding domain (CBD) including the WBM motif (second gate). These trimeric PER1/WDR5/CRY complexes could, in addition to CRY/PER and CRY/PER/CK1δ complexes ^12^, be incorporated into the early repressive complex upon its assembly at circadian promotors.

In contrast to PER1, the PER2 paralogue, due to its low affinity binding to the WDR5 WBM site, is unable to displace RbBP5 from the WBM pocket (Fig. 6b) and is therefore unlikely to impact on the active MLL1 complex. Furthermore, PER2 accumulates later at circadian promotors than PER1 ^7–9^. Hence, PER2 may not co-exist with the active BMAL1/CLOCK associated MLL1 complex at circadian promotors. Moreover, PER2/WDR5 complexes likely accumulate at a later circadian time than PER1/WDR5 complexes. This ordered progression is further supported by our observation that PER1 and PER2 themselves compete for WDR5 binding (Fig. 6a) and by the lower WDR5 affinity of PER2, which would restrict PER2/WDR5 complex formation to the (compared to PER1 delayed) peak concentration phase of PER2.

The distinct WDR5 interactions of PER1 and PER2 also translate into a different interplay with CRY at the transition towards the repressive complex. As PER2 exclusively binds WDR5 via its WBM motif, that is also part of the CRY interface, CRY – due to its much higher affinity - competitively displaces WDR5 even from a preformed PER2/WDR5 complex to form a PER2/CRY complex (Fig. 2,3). Hence, PER2/WDR5 complexes can only be stable in absence of CRY, perhaps within different subnuclear compartments. Alternatively, posttranslational modifications or additional protein interaction networks could influence the interplay between PER2/WDR5- and PER2/CRY interactions in cells. The location of the WBM motif of PER also allows for other interesting regulatory possibilities: In PER2 the WBM motif is located in the vicinity of a Zinc interface that stabilizes the CRY1-PER2 interaction, and nearby Cys412 in the CRY1 lid, which forms a redox sensitive disulfide bond in CRY1 ^42^ that is opened upon PER2 binding ^21^ (Fig. 3b). Hence, the competitive interplay between WDR5-PER- and CRY1-PER interactions could be regulated in the cell by changing redox conditions, which would affect CRY1 disulfide bond formation and free zinc concentrations and thereby the stability of the PER/CRY complex.

Furthermore, PER2 may bind WDR5 together with a second (yet unknown) potentially stabilizing WIN site ligand, exploiting WDR5’s well established function as a scaffold protein ^24^. Likewise, PER1 could - apart from RbBP5 - bind WDR5 together with another (yet unknown) WBM site ligand, that may be more abundant during the repressive phase. A stabilizing WBM site ligand X within a ternary PER1/WDR5/X complex would also explain our unexpected observation, that the N225A mutation in the WDR5 WBM pocket significantly weakens PER1-WDR5 interactions in cells, but not in the purified proteins (Fig. 4c, 5a).

Our data suggest, that WDR5 could be incorporated into the multi-subunit repressive CRY/PER containing complex in the form of PER1/WDR5/CRY trimers or PER(1/2)/WDR5 dimers. Interestingly, pulldown/mass spectrometry experiments so far only identified WDR5 as a component of the repressive PER complex, when PER1 was used as bait ^9^, but not when pulling with PER2 ^7,10,12,13^. Our finding that WDR5 binds PER1 with significantly higher affinity than PER2 provides a plausible explanation for this discrepancy, and suggests, that the lack of WDR5 in PER2-pulldown experiments could be due to the low PER2-WDR5 binding affinity and may not necessarily reflect the absence of WDR5 in these PER complexes. Incorporation of PER2/WDR5 subcomplexes into the multi-subunit early repressive complex however requires that different PER2 molecules engage with either WDR5 or CRY. This could potentially occur in the context of PAS domain mediated PER/PER homodimers ^18,19^. Within the early repressive complex, WDR5 could, through its distinct interactions with PER1 and PER2 and by coordinating additional binding partners, provide specificity and define the stoichiometry of this complex while building or reorganizing it. Thereby, WDR5 could exert its effect on repressive chromatin modifications such as histone H3K9 methylation ^9^ or histone acetylation. Competing interactions of WDR5 and CRY with PER may assist a structured assembly of the early repressive complex but could also help complex disassembly and reorganization at the transition towards the late repressive phase of circadian transcription.

Consistent with our findings on the different WDR5 interactions of PER1 and PER2, which are correlated with distinct interaction networks with activating and repressing complexes, earlier reports described distinct biological phenotypes for PER1 and PER2 overexpression ^43^ or deletion ^14–17^. PER1 deletion leads to shorter circadian periods in mice ^14,15^ as well as a lower circadian amplitude and shortened period in cultured cells ^16^. If PER1 provides an “intermediate step” in the circadian repression process by interacting with WDR5, both a shortened period and reduced amplitude in its absence would make sense: skipping its role eliminates a programmed delay, while interfering with an organized progression in chromatin states. Our results in cells demonstrate phenotypes consistent with this idea from a WDR5 perspective: pharmacological blockade of WDR5 WIN site interactions significantly shortens the circadian period by 2 to 3 h and weakens PER1-WDR5 interactions (Fig. 5), while WDR5 overexpression increases the circadian amplitude (Fig. 1c). In addition to providing a molecular rationale for the ordered transition between circadian activation and circadian repression, our results might also be of therapeutic interest. WDR5 and MLL1 have already proven to be attractive targets for novel cancer therapies, targeting the opening of chromatin in dividing cancer cells ^34,44^. Our results suggest that a similar rationale might be exploited for circadian rhythm disorders.

## Materials & Methods

### Recombinant expression and purification of PER, WDR5, CRY, MLL1 and RbBP5

PER1, PER2, WDR5, RbBP5 and MLL1 proteins were expressed overnight at 18°C in *E.coli* Rosetta (DE3) cells using Terrific Broth (TB) medium. Cells were harvested by centrifugation at 4750xg, resuspended in lysis buffer and lysed with a Branson® sonifier.

Cells expressing GST-WDR5 23-334 (pCoofy3) and GST-PER1 1013-1291 (pGEX6P1) were lysed in 50 mM Tris pH 8.0, 0.2 M NaCl, 5 % Glycerol and 14 mM β-Mercaptoethanol (ß-ME) and centrifuged for 45 min at 40 000g. The supernatant was loaded on a 20 mL GSH-affinity column, which was washed with at least 5 column volumes of lysis buffer. The bound GST fusion protein was cleaved on column overnight by adding 3C-PreScission protease and 7.6 mM MgCl_2_. *Serratia marcescens* (Sm) nuclease was also added to cut remaining DNA/RNA. The cleaved protein was eluted with lysis buffer, concentrated and subjected to size exclusion chromatography (SEC).

Cells expressing His-PER2 1132-1252 (pEC3CKan) ^21^ and His-MBP-hRbBP5 (Isoform 2, pCoofy4) were lysed in 50 mM Tris pH 8.0, 0.2 M NaCl, 5 % Glycerol, 20 mM Imidazole and 1 mM β-ME. Affinity purification was carried out using a 5 mL His-Trap column (GE-Healthcare) with an elution gradient from 20 mM to 500 mM Imidazole. The eluted protein was cleaved overnight with 3C-PreScission protease (His-MBP-hRbBP5) or directly used for SEC (His-PER2). For SEC, affinity purified PER1/2, RbBP5 and WDR5 proteins were concentrated and run on a Superdex S200 16/60 or S200 10/300 SEC column (*GE Life Science*) with a buffer containing 25 mM HEPES pH 7.5, 150 mM NaCl, 5 % Glycerol and 2 mM DTT.

Cells expressing His-GST-hMLL1 3754-3963 (an MLL1 fragment containing the WDR5 binding WIN motif and the adjacent catalytic SET domain) in the pCoofy3 vector were lysed in 50 mM Tris pH 7.5, 500 mM NaCl, 10 % Glycerol, 1 mM β-ME, 5 mM PMSF, 5 mM MgCl_2_ and Sm nuclease and centrifuged for 45 min at 40,000xg. The supernatant was loaded on a 60 mL GST affinity column. After washing steps with lysis buffer and ATP, 1:100 ratio 3C protease was used to cut the His-GST-Tag on column overnight. Cleaved hMLL1 375-3966 protein was eluted with the wash buffer. The protein was concentrated and applied to a Superdex S75 16/60 SEC column (GE Life Science) with a buffer containing 50 mM Tris pH 7.5, 300 mM NaCl, 5 % Glycerol and 1 mM DTT.

His-tagged CRY2-PHR (1-512) protein was recombinantly expressed in *Sf9* insect cells using the baculovirus expression system. The cells were harvested by centrifugation at 500xg for 10min and resuspended in lysis buffer (50 mM Tris pH 7.1, 300 mM NaCl, 10 % Glycerol, 5 mM β-ME, 30 mM Imidazole, Sm Nuclease and 1 tablet of Protease Inhibitor Cocktail (Roche)) followed by mechanical lysis with a Branson® sonifier. The lysate was centrifuged at 20,000xg for 1 hour before loading the supernatant to a His-Trap affinity column (Cytiva). The protein was eluted with a gradient from 20 mM to 500 mM imidazole. The protein fractions were dialysed adding His-tagged 3C protease in a 1:100 molar ratio. The protease and the cleaved His-Tag were removed by reverse Ni-affinity. The tag-free CRY2-PHR protein was further purified via SEC using a Superdex S200 16/60 column (GE Life Science) with a buffer containing 25 mM HEPES pH 7.5, 300 mM NaCl, 5 % Glycerol and 1 mM DTT. CRY1 1-496 (PHR) was purified from insect cells as described before ^21^.

### Site-Directed Mutagenesis

PER1, PER2 and WDR5 mutants were generated by QuikChange site-directed mutagenesis and verified by sequencing. The mutant constructs were expressed and purified to homogeneity essentially as described for the wild-type proteins.

### Analytical Size exclusion chromatography (SEC) of PER/WDR5-, PER/CRY-, WDR5/RbBP5 and PER/WDR5/RbBP5 complexes

For complex formation, 1:3 ratios (His-PER2 or RbBP5 with excess WDR5) or equimolar ratios (PER1/2 with CRY1; PER1 with WDR5; PER1/2 with WDR5 and RbBP5) of the wildtype or mutant proteins were mixed and incubated overnight on ice. For His-PER2 complexes, 3C-protease was added for the overnight incubation to remove the His-Tag of PER2. The proteins were concentrated to 500 µL and run on a Superdex S200 10/300 or S75 10/300 SEC column (GE Life Science) with a buffer containing 25 mM HEPES pH 7.5, 150 mM NaCl, 5 % Glycerol and 2 mM DTT. The individual proteins were run on the same column as a reference. SEC chromatograms were analyzed with the Origin® version v7.5 and the OriginPro version 2023 (OriginLab Corporation, Northampton, MA, USA) software. Protein fractions were analyzed by SDS-PAGE.

### Analytical SEC of PER/WDR5/CRY- and MLL1/WDR5 complexes

For complex formation, proteins were mixed in an equimolar ratio at a final concentration of 90 µM or 100 µM. For PER/CRY/WDR5 complex formation, PER, CRY and WDR5 proteins were mixed in equimolar ratios, incubated for 60 minutes at 4°C and then injected onto an analytical S200 10/300 SEC column. Alternatively, the PER/WDR5 complex was preformed by overnight incubation at 4 °C and then equimolar amounts of CRY were added and incubated for 60 minutes at 4°C before loading onto an analytical S200 10/300 SEC column. To generate an equimolar MLL1/WDR5 complex, we incubated a mixture of MLL1 and WDR5 proteins at 4°C for 2 hours, run an S200 10/300 SEC column and pooled only SEC peak fractions, which contained the equimolar MLL1/WDR5 complex. Afterwards equimolar amounts of PER1 were added to the preformed MLL1/WDR5 complex and incubated for 60 minutes at 4°C before loading onto an analytical S200 10/300 SEC column. The SEC running buffer consisted of 25 mM HEPES pH 7.5, 300 mM NaCl, 5 % Glycerol and 1 mM DDT. Protein fractions were analyzed by SDS-PAGE. SEC chromatograms were analyzed using Microsoft 365® EXCEL and plotted with the OriginPro version 2023 (OriginLab Corporation, Northampton, MA, USA) software.

### Crystallization

Crystals of the WDR5 23-334 N225A protein (10 mg/mL) were obtained at 20°C using a hanging drop setup with a reservoir solution containing 0.1 M HEPES pH 7.5, 50 mM (NH_4_)_2_SO_4_ and 24 % PEG 3350. Crystals belong to space group C222_1_ with 1 molecule per asymmetric unit and cell constants a=78.5 Å, b= 98.13 Å, c=80.06 Å. The crystals were shock frozen in liquid nitrogen using a cryoprotective solution with 15 % PEG 200.

WDR5 23-334 was co-crystallized with a synthetic peptide (Genscript) including residues 1198 to 1211 of PER2 and an N-terminal tyrosine for absorbance at 280 nm (PER2 Y1198-1211). Crystals of the WDR5/PER2 Y1198-1211 complex (12 mg/mL WDR5, 2 mM PER2 peptide) were obtained at 20°C using a hanging drop setup with a reservoir solution containing 0.1 M Tris pH 8.0, 50 mM NaCl and 13% PEG 8000. Crystals belong to space group P4_2_ 2_1_ 2 with unit cell constants a= b = 78.15 Å, c=101.75 Å and 1 complex per asymmetric unit. The crystals were shock frozen in liquid nitrogen with a cryoprotective solution containing 20 % PEG 200.

### Data Collection, Structure Determination, and Refinement

A 1.38 Å data set was collected of a WDR5 N225A crystal at SLS X06SA (Villigen, Switzerland) and a 1.85 Å data set of a WDR5/PER2Y1198-1211 complex crystal at DESY/EMBL PX13 (Hamburg, Germany). All data sets were processed using XDS ^45^ and evaluated using the half set correlation in the highest resolution shell. The data were scaled with AIMLESS and an R_free_ data set consisting of 5% of the data was created. All structures were solved using PHASER ^46^ for molecular replacement with Apo-WDR5 (PDB 2H9L) as search model. Model building and refinement were performed with Coot ^47^ and Refmac ^48^. The models of WDR5 N225A and WDR5/PER2Y1198-1211 show very good stereochemistry and R_work_/R_free_-values of 0.194/0.218 (WDR5 N225A) and 0.1651/0.2007 (WDR5/PER2Y1198-1211).

The WDR5/PER2 Y1198-1211 structure consists of 314 amino acids, where WDR5 23-29 and PER2 Y1198-1202 were not visible in the electron density due to conformational disorder. Additionally, the model includes 98 water molecules, 2 Tris molecules, 1 Polyethylenglycol molecule and 2 Chloride ions. PER2 T1203 was modeled as alanine due to conformational disorder of the side chain. The Ramachandran plot shows 94.19 % of the main chain torsion angles in the most favored regions and no residues marked as outliers.

The WDR5 23-334 N225A structure consists of 301 amino acids, where residues 23-31 and 334 were not visible in the electron density and K227 and R181 (near A225) were modeled as alanine due to conformational disorder. 245 waters are included in the model. The Ramachandran plot shows 94.33 % of the main chain torsion angles in the most favored regions and 0 amino acids marked as outliers. All figures were generated with PyMol.

### Isothermal titration calorimetry

To determine the K_D_ values for the interactions of PER2 1132-1252 and PER1 1013-1291 with either WDR5 23-334 or mCRY1 1-496, and of RbBP5 with WDR5, ITC experiments were performed using a MicroCal PEAQ-ITC Automated system (*Malvern Panalytical, Malvern UK).* All used proteins were dialyzed for at least 16 h to a buffer containing 25 mM HEPES pH 7.5, 150 mM NaCl, 5 % Glycerol and 1mM β-ME. A 750 µM solution of WDR5 (WT or N225A, syringe) was titrated to 50 µM solutions of PER1 1013-1291 (WT, R1111A, D1197A, R1111A D1197A, R1111A R1113A), PER2 1132-1252 (WT, I1205G, D1206A, V1207A) or RbBP5 (cell). For the PER-CRY1 interaction, 90 µM solutions of PER1 1013-1291 (WT and mutants) or PER2 1132-1252 (wt and mutants) (syringe) were titrated into an 8 µM CRY1 1-496 solution (cell). The titration consisted of 19 injections of 2 µl every 150 s at 25 °C. As a control the protein in the syringe was titrated into a cell filled with buffer. The analysis of the data was performed with the Microcal PEAQ-ITC Analysis software, using a one-site binding model. All experiments were performed in triplicates using three distinct samples and a mean of the individual measurements was used for further interpretation.

### Mammalian cell culture and plasmids

WDR5 and WDR5 N225A were cloned into the pEF-dest51 destination vector with a V5 tag (Invitrogen) resulting in the expression clones pEF-dest51-WDR5 and pEF-dest51-WDR5-N225A. Full length PER1 and PER2 were cloned into the pEZYflag vector with a flag tag (Addgene #18700) resulting in the expression clones PER2 WT_pEZYflag, PER2 R634A_pEZYflag, PER2 I1205G_pEZYflag, PER2 D1206A_pEZYflag, PER2 V1207A_pEZYflag, PER1 WT_pEZYflag, PER1 R1111A_R1113A_pEZYflag, PER1 D1197A_pEZYflag, PER1 D1197 R1111_pEZYflag. The final expression clones were used for transient transfection experiments. Additional plasmids used: pWPT-GFP (Addgene #12255), pBMPlucUTR and pcDNA-CRY1V5His ^9,35^.

Human Embryonic Kidney 293 (HEK293) cells and Human Bone Osteosarcoma Epithelial (U2OS) cells were maintained in Dulbecco’s Modified Eagle Medium (DMEM) with high glucose, L-glutamine, and phenol red, supplemented with 1% penicillin/streptomycin (Sigma-Aldrich) and 10% fetal bovine serum (FBS) (Seraglob). HEK293 and U2OS cells were cultured at 37°C in a humidified incubator with 5% CO_2._ They were passaged 1:10 and 1:6, respectively, when they reached about 80% confluence after washing with PBS and detaching with trypsin 1x or 10x, respectively.

### Transient Transfection of HEK293 cells

For Co-IP experiments, HEK293 cells grown to about 50% confluence were transfected using Polyethylenimine (PEI). Each 10 cm cell plate was treated with a mixture of 627 µL 150 mM NaCl, 70 μL PEI5 and 13 μg total DNA. Equal μg amounts of DNA (i.e. 6.5 µg each) were added for the two plasmids used in the Co-IP experiments. The PEI5-DNA mix was incubated 45 minutes at room temperature before being added to the cell culture. The culture medium was changed 24 hours later, and cells were collected 48 hours post transfection.

### Transient Transfection of U2OS cells

For circadian oscillation experiments, U2OS cells were grown to about 80% confluence in black 24-well plates with 0.5 mL culture medium per well. For each well, 50 μL of serum-free medium, 0.5 μg of total plasmid DNA and 1.5 μL of TransIT ®-2020 Transfection Reagent (Mirus Bio, WI, USA) were mixed by gentle pipetting. This mixture was incubated at room temperature for 20 minutes, then added dropwise to the U2OS cells for transfection. For the WDR5 overexpression experiment, where WDR5 was transiently overexpressed along with a BMAL1-luc reporter construct, the relative ratio (weight:weight) of Bmal1-luc (pBMPlucUTR) reporter construct to protein expression construct (pWPT-GFP or pEF-dest51-WDR5) was 1:2.

### Co-Immunoprecipitation (Co-IP) and Western Blotting

HEK293 cells were trypsinized, and for the WDR5-PER pulldown fixed for 3 minutes with 1 % formaldehyde in PBS, then centrifuged for 3 minutes at 1,200xG before quenching with ice cold 1.25 M glycine in PBS. After a PBS wash, fixed cells (WDR5-PER pulldown) or unfixed cells (CRY1-PER pulldown) were lysed with RIPA buffer containing 150 mM NaCl, 50 mM Tris, 1% NP40, 15 mM MgCl2, 5 % glycerol, 1 mM EDTA and protease inhibitors (pepstatin A, leupeptin, aprotinin, trypsin inhibitor, 0.5 mM PMSF). Lysates were sonicated twice for 30 seconds and then centrifuged at 16,000xG for 10 minutes at 4°C. Supernatants were collected and protein concentrations were estimated via Coomassie dot blot relative to an albumin standard. Immunoprecipitation was carried out overnight using an anti-V5-tag antibody from rabbit (LabForce, MBL, 4015-PM003) to precipitate WDR5-V5 or CRY1-V5 or a rabbit IgG antibody as negative control. Unblocked protein A beads (Diagenode, C03020003) were used to pulldown the antibody-protein complexes. The beads were washed three times with lysis buffer before denaturation with SDS sample buffer. Lysates and immunoprecipitates were separated by 10 % SDS-PAGE. Western blots were performed as previously described (Current Protocols in Molecular Biology, Wiley) using 0.1 µm Whatman Protran BA79 Blotting Membrane (Huberlab, Whatman,12.4020.04). A monoclonal anti-flag M2 mouse antibody (Sigma, F1804-200UG) was used to detect the pulled down PER1/2-Flag. An Anti-V5-tag mAb mouse antibody (LabForce AG (MBL), 4015-M167-3) was used to detect precipitated WDR5-V5 or CRY1-V5. A LI-COR® IRDye® 680RD Goat anti-Mouse IgG secondary antibody was used for visualization and blots were imaged with a LI-COR® Odyssey® scanner. Quantification was performed using the Image Studio™ software. After background subtraction, individual band intensities were normalized to the wildtype sample and to overall expression levels in the lysate wherever possible. Where not possible an IP was conducted to concentrate these lysates. For IP blots the ratio between proteins, for example PER2 relative to WDR5, were calculated. This allowed us to quantify how much PER1 or PER2 were pulled down by a given amount of the “bait” protein (WDR5 or CRY1). At least three independent replicates using distinct samples were generated for each experiment.

### Bioluminescence Measurements and Analysis

To assess the involvement of WDR5 in human circadian rhythms, a circadian *in vivo assay* was used, which measures luminescence as a readout of *Bmal1* transcription using a Bmal1-luc (pBMPlucUTR) reporter construct ^9,49^. Transiently transfected U2OS cells were synchronized 24 hours post transfection by a 30-minute incubation with 1 μM dexamethasone, and then washed once with PBS. Next, the assay medium (DMEM without phenol red (Gibco Thermo Fisher) supplemented with glutamax, 10% FBS, 1% penicillin/streptomycin, 10 mM HEPES, and 100 μM luciferin) was added. Plates were transferred to a custom-made bioluminescence reader for bioluminescence recording. Results were analyzed by the Lumicycler Analysis program from Actimetrics (Wilmette, IL, USA). Raw data were detrended, and parameters including period and amplitude were determined by the software as previously described ^50^.

### Application of C6 inhibitor in Co-IP and Bioluminescence experiments

To analyze the effect of the WDR5-IN-4-TFA (C6) compound (MCE, HY-111753A, MEdChemTronica, Sweden) on circadian oscillations, U2OS cells stably infected with a lentiviral Bmal1-luciferase circadian reporter were used ^16^. Cells were synchronized with 1 μM dexamethasone and subsequently incubated with the C6 compound at final concentrations of 0.5 µM, 1 µM, 2 µM, 5 µM, 7.5 µM and 10 µM C6. The C6 compound was dissolved in 100% DMSO and diluted to the applied concentrations with cell culture medium. The final percentage of DMSO did not exceed 0.1% except for the 7.5 µM C6 (0.175% DMSO) and 10 µM C6 (0.2% DMSO) concentrations. To analyze the effect of the C6 compound on PER1-WDR5 interactions in Co-IP experiments, a 5 µM C6 solution was added to the HEK293 cells 24 h post transfection and 24 h prior to cell fixation and collection. No effect of the inhibitor on cell division or viability was observed.

### Statistical Analysis

GraphPad Prism 5 was used for data presentation and statistical analysis (GraphPad Software Inc., California, USA). A Student’s two-tailed t test or two-way ANOVA were used when appropriate and as described in figure legends, *p<0.05, **p<0.01, ***p<0.001. Data are shown as means ± S.E.M. unless otherwise specified.

## Data Availability

Coordinates for the crystal structures of WDR5 N225A (PDB ID 8OK1) and the WDR5 PER2 complex (PDB ID 8OKF) were deposited in the wwPDB database.

## Acknowledgements

We thank Dr. Roberto Orru for his help with crystal structure determination and the staff at SLS and DESY as well as Dr. Andreas Krämer for help with synchrotron data collection. We thank Dr. Christian Kersten for help with ITC measurements and access to the MicroCal ITC device (DFG, GZ: INST 247/921-1 FUGG). We thank the IMB Media laboratory, Protein production and Proteomics Core facilities for continuous support, in particular Dr. Martin Möckel (IMB protein production core facility) for generating the WDR5 triple mutant constructs and helpful discussions. We also thank Ermanno Moriggi for helpful discussions regarding the cellular experiments. Marcel Conrady (MC) and Florian Hof (FH) are supported by the Deutsche Forschungsgemeinschaft (DFG, German Research Foundation), grant numbers GRK2526/1-Project-ID 407023052 (MC) and SFB 1361-Project-ID 393547839 (FH). Steven Brown and Audrey Rawleigh were financed by a grant from the Swiss National Science Foundation, as well as additional support from the Molecular Life Sciences graduate program and the Human Frontiers Science Program.

Finally, we would like to express our deepest sorrow about the sudden death of our highly appreciated collaborator and supervisor Steven Brown.

## Extended Data

**Extended Data Table 1:** X-ray data collection and refinement statistics of WDR5 N225A and WDR5/PER2 Y1198-1211.

|  | WDR5 23-334 N225A | WDR5 23-334/PER2 Y1198-1211 |
| --- | --- | --- |
| *Data collection* |  |  |
| **Beamline** | SLS X06SA | DESY PX13 |
| **Wavelength (Å)** | 0.9999 | 0.9763 |
| **Resolution range (Å)** | 48.67–1.38 (1.40–1.38) | 78.15–1.85 (1.89–1.85) |
| **Space group** | C 2 2 21 | P 42 21 2 |
| **a, b, c (Å)** | 78.498, 98.134, 80.055 | 78.152, 78.152, 101.748 |
| **$\alpha$, $\beta$, $\gamma$ (°)** | 90, 90, 90 | 90, 90, 90 |
| **Unique reflections** | 63526 (2949) | 27636 (1653) |
| **Multiplicity** | 1.9 (2.0) | 14.2 (14.4) |
| **Completeness (%)** | 99.3 (93.1) | 100 (100) |
| **Mean I/$\sigma$(I)** | 13.0 (1.2) | 18.3 (5.2) |
| **CC<sub>1/2</sub>** | 0.999 (0.573) | 0.999 (0.955) |
| **R<sub>merge</sub>** | 0.028 (0.291) | 0.103 (0.541) |
| *Refinement* |  |  |
| **Protein atoms** | 2349 | 2406 |
| **Water molecules** | 245 | 98 |
| **R<sub>work</sub>** | 0.1944 | 0.1651 |
| **R<sub>free</sub>** | 0.2177 | 0.2007 |
| **RMSD bonds (Å)** | 0.013 | 0.012 |
| **RMSD angles (°)** | 1.802 | 1.647 |
| **B-factors overall** | 19.43 | 20.15 |
| **B-factors** | 18.26 (WDR5), 30.74 (H<sub>2</sub>O) | 19.53 (WDR5), 32.13 (PER2), 23.94 (H<sub>2</sub>O) |
| **Favored (%)** | 94.33 | 94.19 |
| **Allowed (%)** | 5.67 | 5.81 |
| **Outliers (%)** | 0 | 0 |

**Extended Data Table 2:** ITC results of PER1 1013-1291, PER2 1132-1252, WDR5 23-334, CRY1 1-496 and RbBP5 (WT and mutants) protein interactions. Average values of three measurements with standard deviations are shown.

| Cell / Analyte | Syringe / Titrant | N (sites) | $K_D$ (μM) | $\Delta H$ (kJ/mol) | $\Delta G$ (kJ/mol) | $-T\Delta S$ (kJ/mol) |
| --- | --- | --- | --- | --- | --- | --- |
| **PER1 WT** | **WDR5 WT** | $0.8 \pm 0.003$ | $3.09 \pm 0.20$ | $-46.13 \pm 0.62$ | $-31.5 \pm 0.17$ | $14.70 \pm 0.77$ |
| PER1 R1111A | WDR5 WT | $1.0 \pm 0.01$ | $13.47 \pm 1.48$ | $-36.80 \pm 1.58$ | $-27.87 \pm 2.99$ | $8.93 \pm 1.86$ |
| PER1 D1197A | WDR5 WT | $1.8 \pm 0.007$ | $5.65 \pm 0.16$ | $-45.87 \pm 0.48$ | $-30.00 \pm 0.06$ | $15.9 \pm 0.54$ |
| PER1 R1111A D1197A | WDR5 WT | $0.8 \pm 0.03$ | $16.17 \pm 1.71$ | $-44.10 \pm 3.29$ | $-27.40 \pm 0.28$ | $16.67 \pm 3.55$ |
| PER1 R1111A R1113A | WDR5 WT | Weak binding* | $> 100$ | | | |
| PER1 WT | WDR5 N225A | $0.8 \pm 0.01$ | $7.26 \pm 0.24$ | $-35.77 \pm 0.61$ | $-29.4 \pm 0.096$ | $6.38 \pm 0.7$ |
| **PER2 WT** | **WDR5 WT** | $0.9 \pm 0.1$ | $50.73 \pm 19.33$ | $-16.87 \pm 5.195$ | $-24.93 \pm 0.91$ | $-7.99 \pm 6.08$ |
| PER2 I1205G | WDR5 WT | Not binding |  |  |  |  |
| PER2 D1206A | WDR5 WT | Not binding |  |  |  |  |
| PER2 V1207A | WDR5 WT | Weak binding* | $69.91 \pm 36.90$ | | | |
| PER2 WT | WDR5 N225A | Not binding |  |  |  |  |
| **CRY1** | **PER1 WT** | $0.8 \pm 0.004$ | $0.016 \pm 0.006$ | $-87.70 \pm 3.23$ | $-44.80 \pm 1.02$ | $42.87 \pm 4.25$ |
| CRY1 | PER1 R1111A | $0.9 \pm 0.02$ | $0.0098 \pm 0.0035$ | $-76.27 \pm 0.73$ | $-46.03 \pm 0.86$ | $30.3 \pm 0.48$ |
| CRY1 | PER1 D1197A | $1.0 \pm 0.02$ | $0.157 \pm 0.036$ | $-66.53 \pm 1.64$ | $-38.97 \pm 0.53$ | $27.53 \pm 2.00$ |
| CRY1 | PER1 R1111A D1197A | $1.0 \pm 0.006$ | $0.196 \pm 0.032$ | $-63.60 \pm 0.71$ | $-38.40 \pm 0.415$ | $25.23 \pm 1.08$ |
| **CRY1** | **PER2 WT** | $0.9 \pm 0.05$ | $0.035 \pm 0.013$ | $-54.38 \pm 5.41$ | $-43.28 \pm 1.15$ | $11.10 \pm 6.06$ |
| CRY1 | PER2 I1205G | $0.8 \pm 0.01$ | $0.086 \pm 0.012$ | $-80.25 \pm 2.16$ | $-40.43 \pm 0.31$ | $39.83 \pm 2.20$ |
| CRY1 | PER2 D1206A | $0.8 \pm 0.04$ | $0.069 \pm 0.025$ | $-46.93 \pm 3.41$ | $-41.48 \pm 0.95$ | $5.45 \pm 3.86$ |
| CRY1 | PER2 V1207A | $0.9 \pm 0.06$ | $0.037 \pm 0.013$ | $-63.73 \pm 0.75$ | $-48.53 \pm 6.70$ | $15.20 \pm 7.20$ |
| **RbBP5** | **WDR5 WT** | $1.0 \pm 0.003$ | $0.61 \pm 0.014$ | $-35.97 \pm 0.22$ | $-35.50 \pm 0.063$ | $0.48 \pm 0.27$ |
| RbBP5 | WDR5 N225A | Weak binding<br>$1.6 \pm 0.1$ | $71.00 \pm 18.12$ | $-9.09 \pm 1.71$ | $-23.87 \pm 0.67$ | $-14.73 \pm 2.31$ |

\* N, $\Delta H$ , $\Delta G$ and $-T\Delta S$ were omitted due to very weak binding.

**Extended Data Figure 1:**
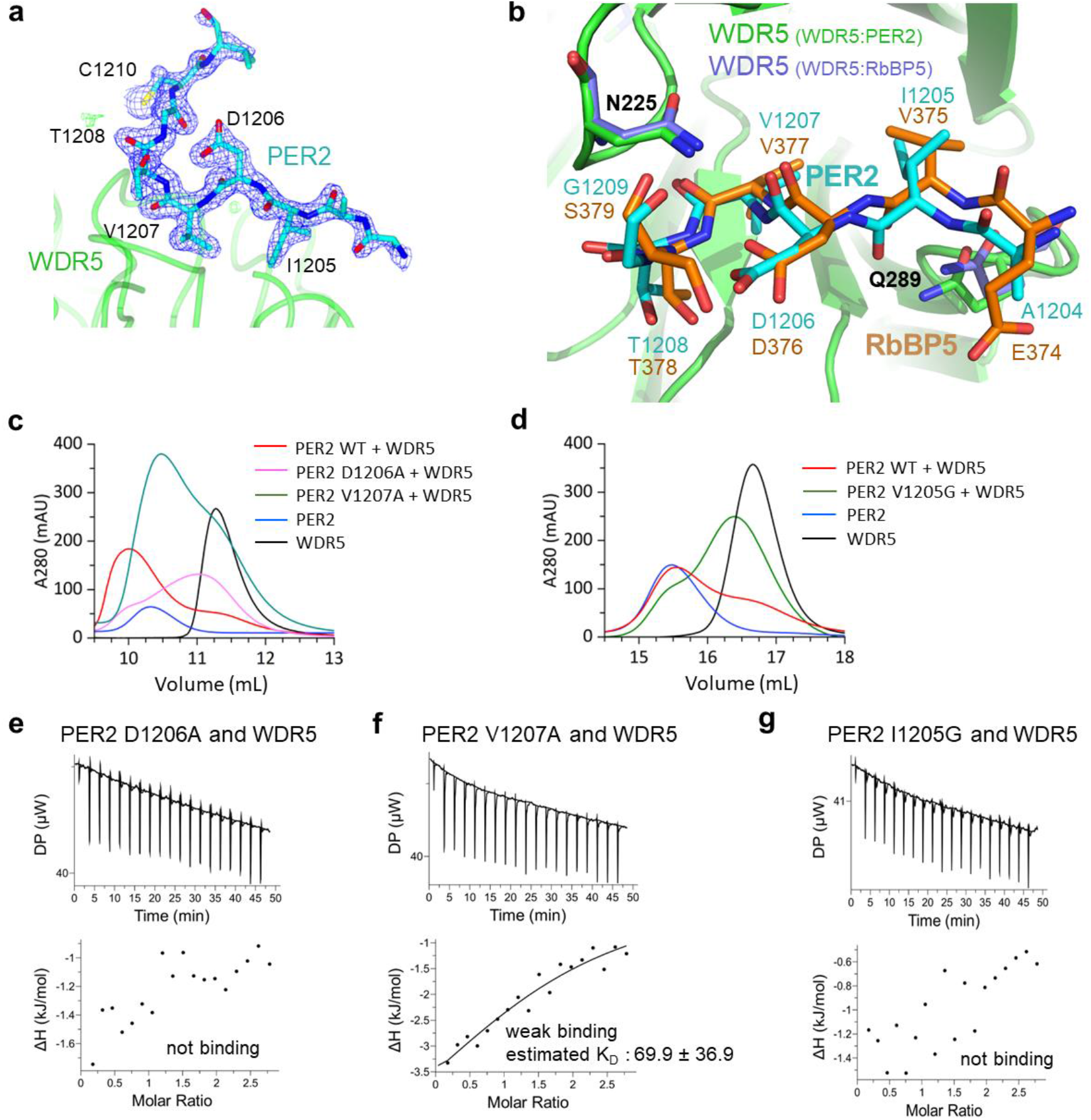
Interaction of PER2 with WDR5. **a** 2Fo-Fc electron density map of the WDR5-bound PER2 WBM motif peptide (Y^1198^TGGLPTAIDVTGCV^1211^) at 1.35σ (blue). Residues T1203 to V1211 of the PER2 peptide (cyan) are well resolved in the electron density. T1203 is modelled as alanine. WDR5 is shown in green. **b** Close-up view of the WBM motif of RbBP5 (orange, PDB 2XL3) ^29^ and PER2 (cyan) binding to WDR5. WDR5 of the WDR5/PER2 complex is shown in green. N225 and Q289 of the WDR5/RbBP5 complex are shown in purple. The structures were superimposed on WDR5 with an RMSD of 0.271 Å **c,d** SEC chromatograms of WDR5 23-334 and PER2 1132-1252 (WT, I1205G, D1206A, V1207A). **c** Chromatograms of excess WDR5 mixed 3:1 with PER2 WT (red), PER2 D1206A (magenta) or PER2 V1207A (green). WDR5 23-334 alone (black) and PER2 1132-1252 WT alone (blue) are shown for comparison. An S75 10/300 column was used. **d** Chromatograms of excess WDR5 mixed 3:1 with PER2 WT (red) or PER2 I1205G (green). WDR5 23-334 alone (black) and PER2 1132-1252 WT alone (blue) are shown for comparison. An S200 10/300 column was used. **e,f,g** Representative ITC experiments for **e** PER2 1132-1252 D1206A, **f** V1207A and **g** I1205G with WDR5 23-334 as titrant are shown. The K_D_-value for PER2 1132-1252 V1207A (f) is the means of the triplicates +/- SD, also shown in Extended Data Table 2 and Fig. 2e. *Upper panel*: raw data, *lower panel*: integrated heat (points) with fit for a one-site binding model (line). For PER2 1132-1252 D1206A (e) and I1205G (g) no binding was observed.

**Extended Data Figure 2:**
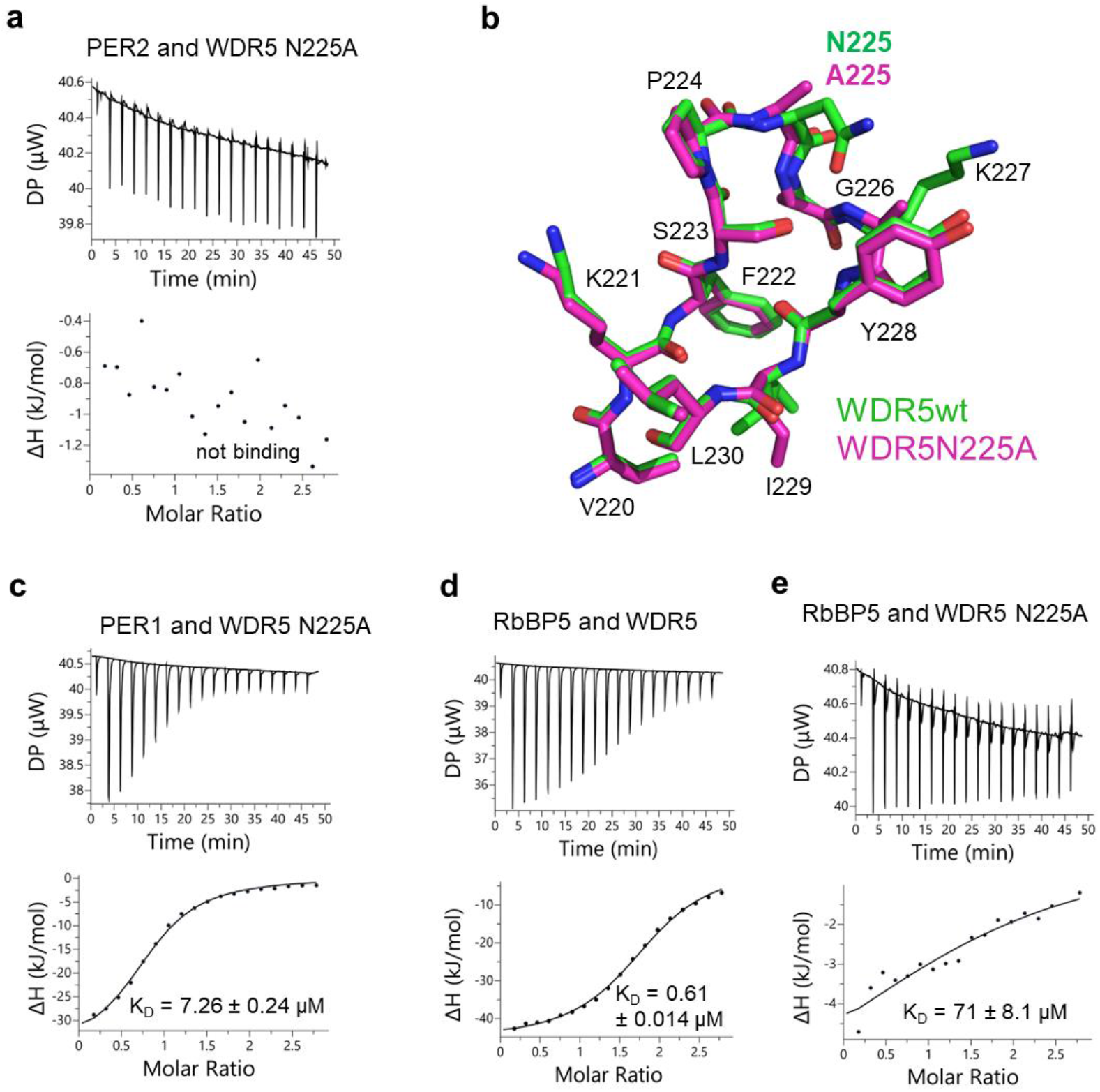
Impact of WDR5 N225A mutation on PER2-WDR5-, PER1-WDR5- and RbBP5-WDR5 interaction. **a,c,d,e** A representative ITC experiment for PER2 1132-1252 with **a** WDR5 23-334 N225A, **c** PER1 1013-1291 with WDR5 23-334 N225A, **d** RbBP5 with WDR5 23-334 WT, and **e** RbBP5 with WDR5 23-334 N225A is shown. The K_D_ values are the means of the triplicates with standard deviations (+/- SD), also shown in Extended Data Table 2 and Fig. 4c. *Upper panel*: raw data, *lower panel*: integrated heat (points) with fit for a one-site binding model (line). For PER2 1132-1252 with WDR5 N225A (a) no binding was observed (Extended Data Table 2, Fig. 2e). **b** Close-up view of the WDR5 N225A mutant crystal structure (magenta) superimposed with the WDR5 WT structure (green, PDB 2H9L) ^51^ around amino acid N225 (RMSD=0.343 Å). K227 is modeled as alanine in the WDR5 N225A structure due to conformational disorder.

**Extended Data Figure 3:**
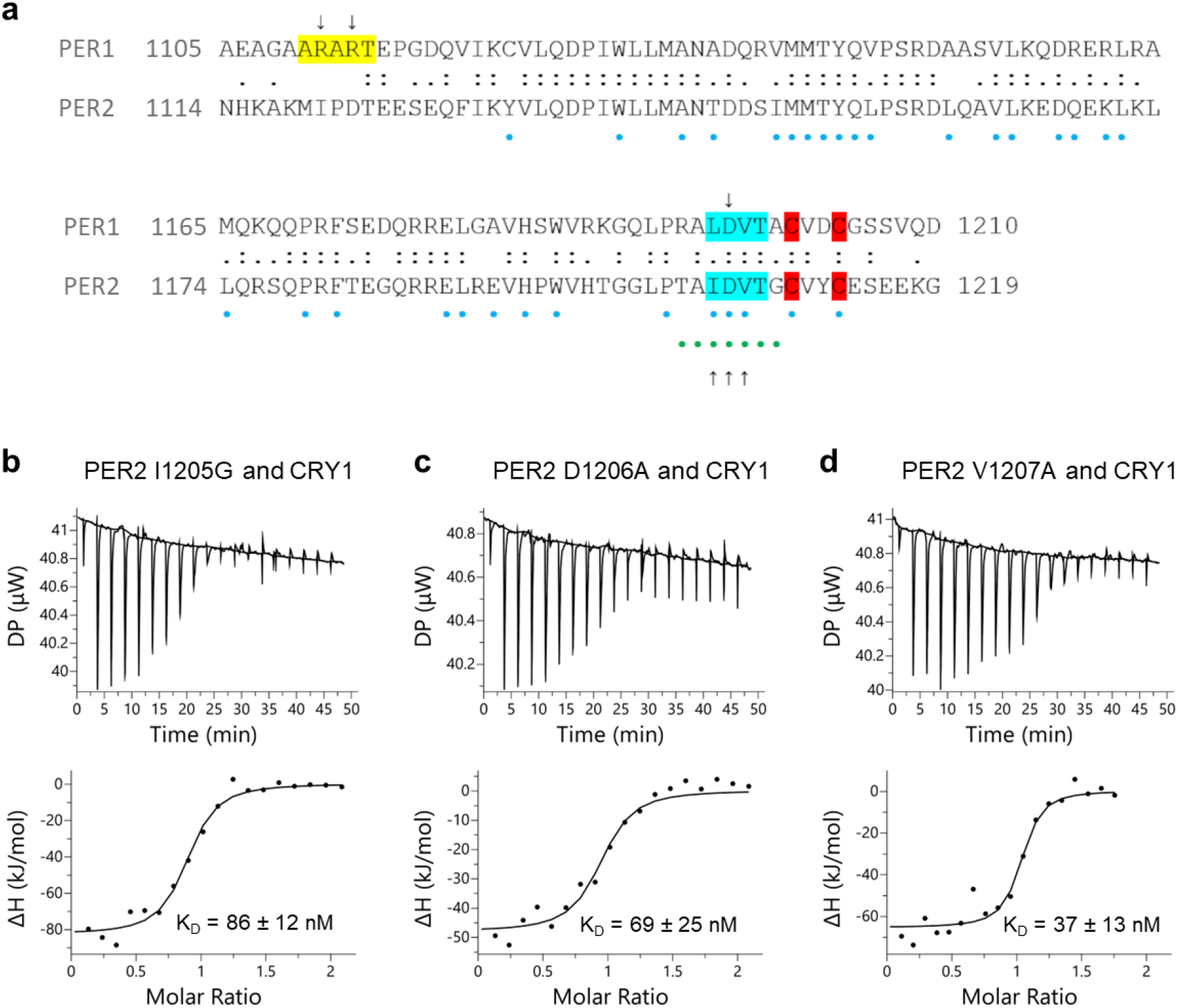
Sequence alignment of PER1 and PER2 and interaction of PER2 1132-1252 with CRY1. **a** Alignment of PER1 1105-1210 and PER2 1114-1219. Identical amino acids are shown by “:” and similar amino acids by “.”. The WDR5 binding motifs are shown in cyan (WBM) and yellow (WIN). Important amino acids of PER2 for interaction with CRY1 (blue) (Schmalen *et al,* 2014) or WDR5 (green) (Fig. 2) are highlighted as dots. Amino acids mutated within this study are indicated by arrows under (PER2) or above (PER1) the alignment. Amino acids that interact with CRY1 or with the WDR5 WBM site are mostly conserved between PER1 and PER2. The WIN motif of PER1 is not conserved in PER2. **b-d** A representative ITC experiment for **b** PER2 1132-1252 I1205G, **c** PER2 D1206A and **d** PER2 V1207A (titrant) with CRY1 1-496 (PHR) is shown. The K_D_ values are the means of the triplicates +/- SD, also shown in Extended Data Table 2 and Fig. 3c. *Upper panel*: raw data, *lower panel*: integrated heat (points) with fit for a one-site binding model (line).

**Extended Data Figure 4:**
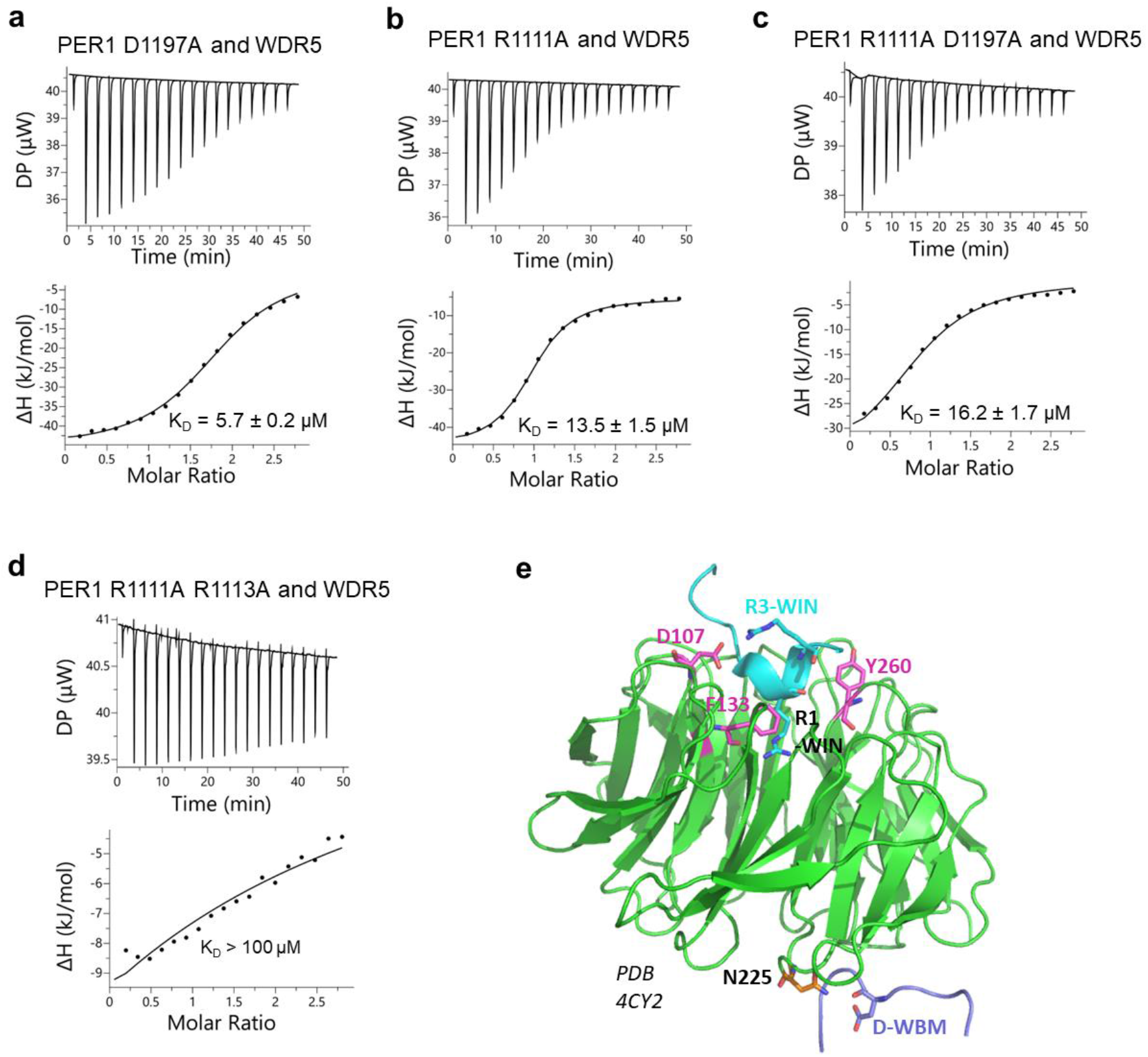
Interaction of PER1 1013-1291 with WDR5. **a-d** A representative ITC experiment for **a** PER1 1013-1291 D1197A, **b** PER1 R1111A, **c** PER1 R1111A D1197A and **d** PER1 R1111A R1113A with WDR5 23-334 (titrant) is shown. The K_D_ values are the means of the triplicates +/- SD, also shown in Extended Data Table 2 and Fig. 4c. *Upper panel*: raw data, *lower panel*: integrated heat (points) with fit for a one-site binding model (line). For PER1 R1111A R1113A and WDR5 (d) only very weak binding was observed, with an estimated K_D_ > 100 µM. **e** Overview of the structure of WDR5 (green) bound to a KANSL1 WIN motif peptide (cyan) in the WIN motif binding pocket and to a KANSL2 WBM motif peptide (purple) in the WBM motif binding pocket of WDR5 (PDB 4CY2) ^36^. Important amino acids in the WIN motif binding pocket of WDR5 (D107, F133, Y260) are shown in pink. The two WIN motif arginine residues of KANSL1 (denoted R1-WIN, R3-WIN) correspond to R1111 (R1) and R1113 (R3) of PER1. N225 in the WBM motif binding pocket of WDR5 is highlighted in orange. The conserved aspartate (denoted D-WBM) in the KANSL2 WBM motif peptide corresponds to D1197 in PER1 and D1206 in PER2.

**Extended Data Figure 5:**
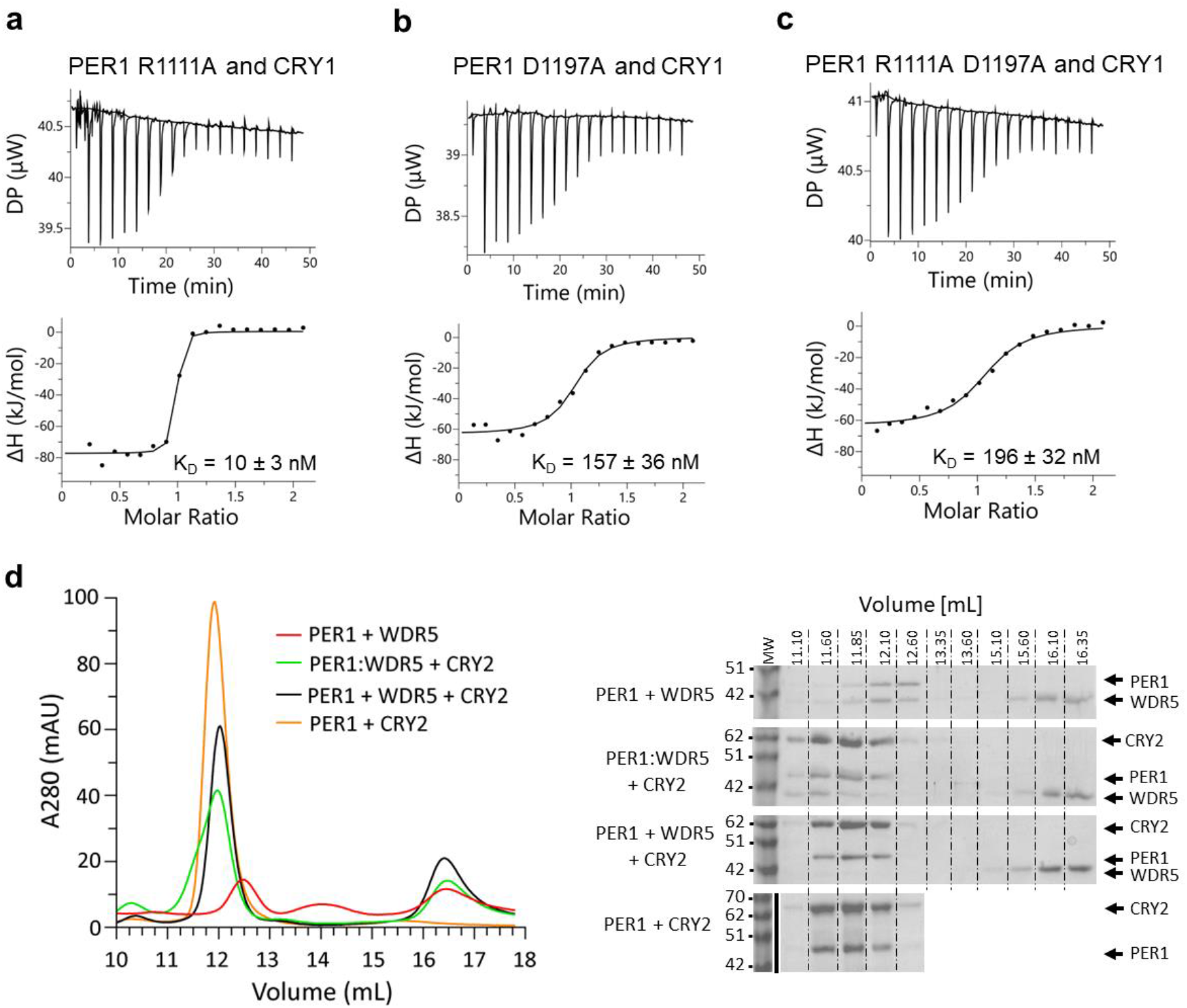
Interaction of PER1 1013-1291 with CRY1 and CRY2. **a-c** A representative ITC experiment for **a** PER1 1013-1291 R1111A, **b** PER1 D1197A and **c** PER1 R1111A D1197A (titrant) with CRY1 1-496 (PHR) is shown. The K_D_ values are the means of the triplicates +/- SD, also shown in Extended Data Table 2 and Fig. 4c. *Upper panel*: raw data, *lower panel*: integrated heat (points) with fit for a one-site binding model (line). **d** Chromatograms (left) and SDS-PAGE analyses (right) of SEC (S200 10/300) of the PER1 1013-1291 interaction with CRY2 1-512 (PHR) and WDR5 23-334. *SDS-PAGE top:* PER1/WDR5 complex (red chromatogram); *second:* preformed PER1/WDR5 complex incubated with equimolar CRY2 (light green chromatogram). A trimeric PER1/WDR5/CRY2 complex is formed, that starts to elute at 11.1 mL; *third:* equimolar mixture of PER1, CRY2 and WDR5 (dark green chromatogram). Only a dimeric PER1/CRY2 complex is observed, WDR5 is displaced; *bottom:* PER1/CRY2 complex (orange chromatogram), that starts to elute at 11.6 mL, i.e. later than the trimeric PER1/WDR5/CRY2 complex.

**Extended Data Fig. 6:**
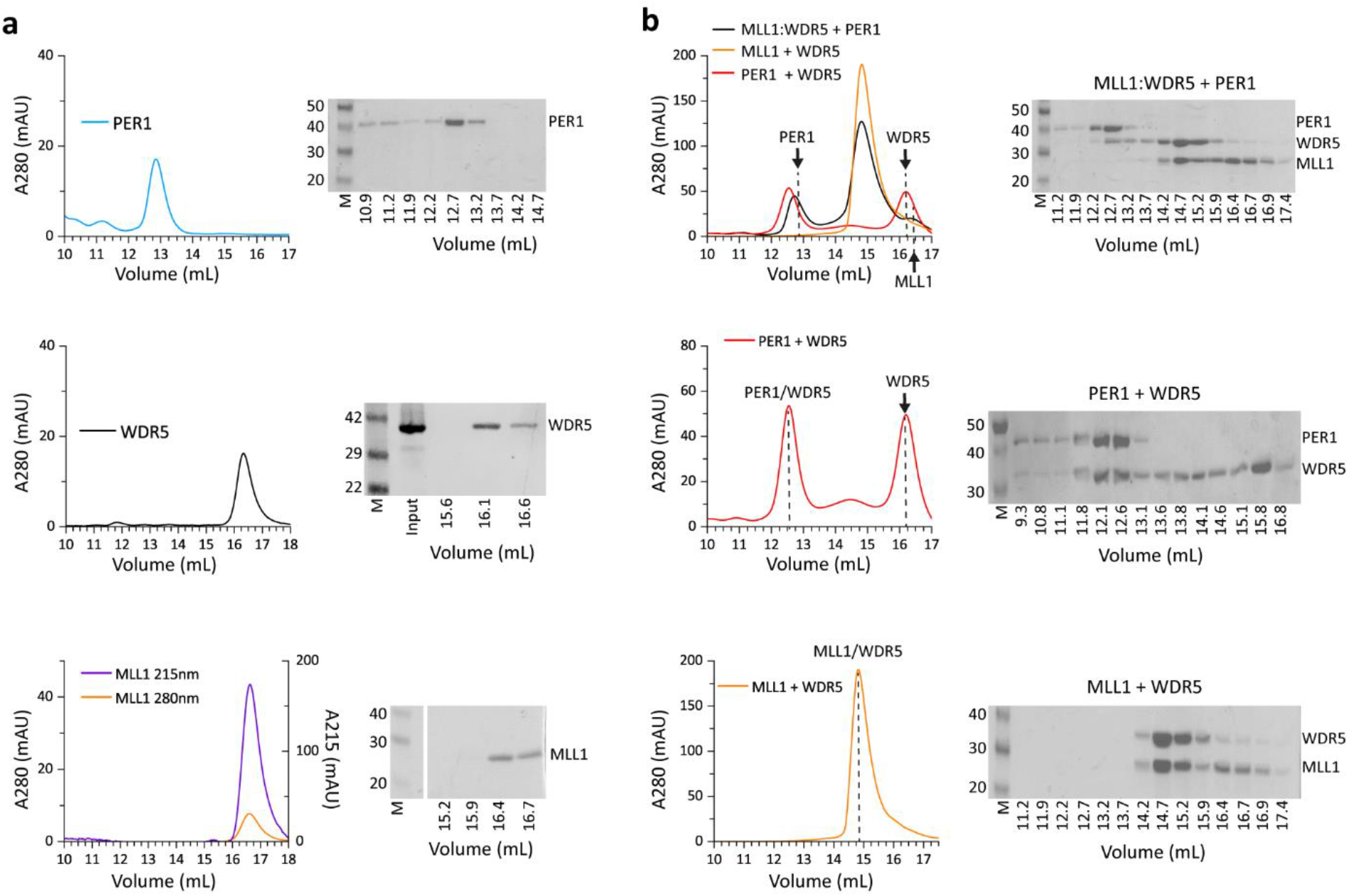
Interaction interplay of PER1 1013-1291 and MLL1 3754-3963 on WDR5. **a** Chromatograms (left) and SDS-PAGE analyses (right) of the SEC (S200 10/300) runs with the individual proteins PER1 1013-1291 (top), WDR5 23-334 (middle) and MLL1 3754 - 3966 (WIN motif + SET domain) (bottom). As MLL1 has a very low absorbance at 280 nm, we also show the chromatogram for absorbance at 215 nm. **b** Chromatograms (left) and SDS-PAGE analyses (right) of SEC (S200 10/300) runs with dimeric PER1/WDR5 (middle) and MLL1/WDR5 (bottom) complexes. *Top*: preformed MLL1/WDR5 complex incubated with equimolar PER1 (black chromatogram). PER1 partially displaces MLL1 from the preformed MLL1/WDR5 complex (identical to Fig. 6d, added for direct comparison with SEC runs of dimeric PER1/WDR5- and MLL1/WDR5 complexes and individual PER1, MLL1 and WDR5 proteins)

